# From Signal to Symphony: Exploring 2D Sequence Representations for Protein Function Prediction

**DOI:** 10.1101/2025.10.26.684638

**Authors:** Yiquan Wang, Minnuo Cai, Yuhua Dong, Yahui Ma, Kai Wei

**Affiliations:** Xinjiang Key Laboratory of Biological Resources and Genetic Engineering, College of Life Science and Technology, Xinjiang University, Urumqi 830049, Xinjiang, China; College of Mathematics and System Science, Xinjiang University, Urumqi 830046, Xinjiang, China; Advanced Research Institute of Multidisciplinary Sciences, Beijing Institute of Technology, Beijing 100081, China

**Keywords:** Protein Function Prediction, Sonification, Biological Signal Processing, Sequence Representation, Generative Protein Design

## Abstract

Predicting protein function from its primary sequence is a fundamental challenge in computational biology. While deep learning has excelled, the optimal representation of sequence data remains an open question. This study explores protein sonification—the conversion of amino acid sequences into 2D spectrograms—as a representation for this task. To facilitate this investigation, we developed a benchmark dataset of 18,000 sequences spanning 12 functionally diverse protein classes. Our systematic evaluation suggests that the structural transformation from a 1D sequence to a 2D spectrogram may be a key contributor to the model’s predictive performance. This observation is supported by ablation studies where models using either purely visual or acoustic features from the spectrogram demonstrated effective standalone performance, suggesting that the representation itself is a key source of this capability. For instance, a model using a sonification map without explicit biophysical meaning achieved 81.08% accuracy, while our biophysically-informed model reached 84.00%, indicating that such domain knowledge may offer a modest performance benefit. When trained from scratch on our dataset, our fusion model achieved performance comparable to or slightly exceeding that of standard transformer architectures like ESM-2 and ProtBERT, suggesting its potential for data efficiency in this specific context. The model’s potential for generalizability was further supported by its performance on the external CARE enzyme classification benchmark, where it achieved 90.44% accuracy. Finally, as a proof-of-concept, we explore the utility of our encoding to guide a diffusion model in generating novel GFP variants, which were assessed for structural viability using computational methods. Our work provides evidence suggesting that the utility of sonification in this context may stem largely from its representational structure, offering a perspective on feature engineering for biological sequences.

## Introduction

Understanding the complex relationship between protein sequence, structure, and function is a paramount challenge in the biological sciences [1–4]. While dominant computational methods, such as Protein Language Models (PLMs) and structure-based predictors, have achieved remarkable success, they each have inherent limitations [5–7]. PLMs, which process sequences linearly, can struggle to capture the long-range, quasi-periodic correlations that define global protein architecture [8–10]. This creates an opportunity for novel data representations that can efficiently extract holistic, functional information from primary sequences.

This study investigates protein sonification, the translation of sequence data into a rich, two-dimensional spectrogram [11–13]. Our approach leverages a powerful and broadly validated computational strategy: transforming 1D biological sequences into 2D representations to capture long-range interactions (LRIs) that are otherwise missed. This paradigm has become a pillar of modern bioinformatics, driving breakthroughs in fields as diverse as protein structure prediction with AlphaFold [14, 15], 3D genome folding [16, 17], and RNA secondary structure analysis [18]. Motivated by these successes, we employ sonification to “fold” a 1D sequence into a 2D spectrogram. In this format, sequence-distant residues are brought into spatial proximity, enabling standard convolutional networks to model global functional signatures in a computationally efficient manner.

However, a critical question arises: if such a method is effective, what is the source of its power? Is it the specific, biophysically-inspired rules used for the translation, which could be viewed as speculative, or is it the fundamental act of transforming a 1D sequence into a 2D representation? To address this question directly, we moved beyond heuristic mappings to establish a quantitative sonification framework based on first principles of amino acid physicochemical properties. We began by rigorously validating the core premise of this framework—the link between static sequence properties and true protein dynamics—with extensive molecular dynamics simulations across diverse protein fold families. Then, we conducted a series of controlled experiments on a broad and functionally diverse dataset, comparing our theory-driven model against alternatives with inverted or randomized semantics. This investigation points towards the importance of the 1D-to-2D data representation itself, a central theme of this work. Our results suggest that functional patterns might be learned as emergent properties from this structured representation.

## Related Works

### Computational Approaches to Protein Function Prediction

The analysis of protein structure and function has long been propelled by experimental techniques such as X-ray crystallography and Cryo-EM, which provide high-resolution but often static snapshots of molecular machinery [19, 20]. To overcome the throughput and dynamic limitations of these methods, computational approaches have become indispensable [21]. In recent years, deep learning has revolutionized the field, most notably with DeepMind’s AlphaFold, which has achieved unprecedented accuracy in predicting protein structures from sequence [7, 22, 23].These advances have driven new biological insights by providing reliable structural models for millions of proteins [24].

However, a static structure does not equate to the complete function. Even state-of-the-art models face challenges in capturing the dynamics of flexible regions, modeling complex multi-protein interactions, and fully elucidating the functional context from structure alone [25, 26].This highlights a critical gap: there is an urgent need for novel data representations that can transcend 1D sequences and 3D structures to more comprehensively capture functional information. While models operating on 1D sequences (like protein language models [27]) or 3D structures [7,15,28]) have been highly successful, there remains a need for novel data representations that can capture the hierarchical and quasi-structural information implicitly encoded in a protein’s primary sequence in a more holistic manner [29].

### Sonification and Audio-Based Methods in Bioinformatics

One emerging avenue for developing such representations is to draw an analogy between the hierarchical complexity of proteins and music, whose mathematical and structural properties offer a rich framework for encoding biological data [30, 31]. This concept was established in early pioneering works like ‘Protein Music’ [32], which translated sequences into melody and bass lines for data analysis. The approach has since been adapted for diverse goals, including enhancing accessibility for the visually impaired by creating classical music from sequences [33, 34] and assisting in complex tasks like protein alignment through combined auditory and visual representations [35].

Building on this foundation, recent work has employed more sophisticated deep learning methods. A significant advancement is Buehler’s ‘AttentionCrossTranslation’ model, which introduced a powerful framework for unsupervised, bidirectional translation between musical and protein domains using interacting transformer networks [12]. A key strength of their model is its ability to perform bidirectional and cycle-consistent translations, ensuring high fidelity. This work established a powerful, generalizable methodology for finding hidden relationships between data types, moving the field towards fully automated pattern discovery. Concurrently, the “conversion of music to protein” (CoMtP) concept further demonstrated that musical scores can serve as templates for designing new functional peptides [36].

While the unsupervised discovery of translation rules, as pioneered by Buehler et al., is a powerful approach for fundamental pattern analysis, our research addresses a different, complementary scientific goal. Our work is not aimed at discovering a universal mapping itself, but rather at testing a specific, human-driven hypothesis: can a biophysically-grounded, quantitative mapping serve as an effective feature engineering strategy for a downstream machine learning task, namely protein function prediction? Therefore, our philosophy is distinct. We intentionally leverage existing scientific knowledge (i.e., physicochemical properties) to construct a transparent and interpretable quantitative mapping. The primary goal is not translation fidelity for its own sake, but the creation of a feature-rich 2D representation (the spectrogram) that is optimized for analysis by computer vision models. The effectiveness of this knowledge-driven, feature engineering approach is evaluated by the accuracy achieved on the predictive task.

### The Resonant Recognition Model: A Precedent for Signal-Based Functional Analysis

A significant precedent for treating biological sequences as signals to decode function is the Resonant Recognition Model (RRM). This physico-mathematical framework posits that biomolecular interactions are governed by resonant energy transfer [37, 38]. The core methodology converts an amino acid sequence into a numerical signal—often using the electron-ion interaction potential (EIIP)—and then applies Fourier analysis. This approach builds directly on foundational work that first demonstrated how periodicities in physicochemical properties, such as the “hydrophobic moment,” strongly correlate with secondary structures like *α*-helices and *β*-sheets [39]. The RRM posits that a single, dominant peak in the resulting frequency spectrum—the “characteristic frequency”—is strongly correlated with the protein’s specific biological function or interaction [40]. The practical utility of this model is well-established; it has been successfully used to design novel bioactive peptides [41] and, more recently, to explain complex phenomena like long-distance DNA-protein interactions [42].

Furthermore, the RRM framework has proven to be both generalizable and extensible. Its principles have been successfully adapted for the classification of other macromolecules, serving as an effective preprocessing technique for distinguishing between different classes of RNA sequences [43]. Recognizing that a single power spectrum might not capture all available information, researchers developed the Complex RRM (CRRM). This extension utilizes both the real and imaginary components of the Fourier spectrum, providing a richer, two-dimensional view of the signal’s frequency characteristics. The CRRM demonstrated superior performance in distinguishing subtle biological differences between closely related viral protein families, such as subtypes of Influenza A neuraminidase, where the traditional RRM was insufficient [44].

Collectively, the extensive literature on the RRM and its derivatives provides a powerful theoretical and empirical foundation for our work. It establishes that transforming the linear information of a biological sequence into the frequency domain is a biophysically-grounded and highly effective strategy for functional analysis. The evolution from RRM to CRRM also highlights a clear trajectory: increasing the dimensionality of the signal representation can unlock deeper biological insights. Our sonification approach is conceptually aligned with this trajectory but takes a substantial leap forward by synthesizing multiple physicochemical properties into a complex, information-dense two-dimensional spectrogram.

## Methods

### Dataset Curation and Preprocessing

To construct a benchmark for evaluating protein function prediction that represents a broad spectrum of biological roles, we curated a dataset of 18,000 sequences from twelve distinct protein classes: Enzyme, Structural, Transport, Storage, Signalling, Receptor, Regulatory, Immune, Chaperone, Cell Adhesion, Motor, and Antimicrobial. Protein sequences were sourced from several authoritative public databases to ensure diversity and quality. These included the NCBI RefSeq collection [45], the UniProt Knowledgebase (UniProtKB) [46], and specialized databases such as the Antimicrobial Peptide Database (APD) [47], the Transporter Classification Database (TCDB) [48] for transport proteins, and the CARE database [49] for enzymes. To ensure data quality and reduce redundancy, we employed the CD-HIT clustering algorithm [50], removing all sequences with an identity of over 90% to any other sequence in the dataset. The final dataset was balanced, containing 1,500 sequences for each of the twelve classes. For model training and evaluation, we partitioned the data into a training set (70%, 12,600 sequences), a validation set (10%, 1,800 sequences), and an independent test set (20%, 3,600 sequences). This stratified split ensures that each data partition maintains the original class distribution [51].

### Quantitative, Biophysically-Grounded Sonification Framework

To move beyond heuristic rules and establish a rigorous foundation, we developed a quantitative framework that maps fundamental amino acid physicochemical properties to musical elements. The core of our framework is the translation of the primary sequence into a musical score, which is then rendered into a 2D spectrogram for analysis. Our central scientific question was whether the model’s performance stems from the specific biophysical semantics encoded in the mapping rules, or from the structural transformation of the 1D sequence into a 2D representation. To answer this, we implemented and compared three distinct mapping schemes.

#### Theory-Driven Quantitative Model

This model serves as our principled, theory-driven starting point. Our focus on hydrophobicity is motivated by the foundational discovery that periodicities in this property along the primary sequence are strong determinants of secondary structure [39], with further computational studies confirming that this global patterning of hydrophobic and polar residues is a dominant force in determining the overall protein fold, often overriding the intrinsic propensities of individual amino acids [52] Building on this, our model is based on the biophysical hypothesis that stable, hydrophobic residues in the protein core correspond to low-frequency collective motions. To implement this, we map higher hydrophobicity to lower pitches, a design inspired by psychoacoustic analogies linking low frequencies to stability [53]. The validity of this choice is empirically demonstrated in our results (Table 2), where this mapping provides a clear performance advantage over control models. The pitch of each note is determined by the hydrophobicity of the corresponding amino acid, following the formula:

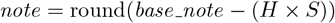

where `note` is the resulting MIDI pitch number, `base_note` is a reference pitch set to 60 (Middle C), `H` is the Kyte-Doolittle hydrophobicity index of the amino acid, and `S` is a scaling factor set to 3. This formula directly implements our hypothesis that higher hydrophobicity corresponds to lower, more stable-sounding pitches. Timbre, or the tonal quality, is determined by amino acid polarity based on the Zimmerman scale, with polar residues mapped to “bright” timbres (e.g., piano) and non-polar residues to “dark” timbres (e.g., cello).

#### Control Models

To deconstruct the source of performance, we designed two control models:

1. **Inverted Semantics Model:** As a direct control for the directionality of the biophysical analogy, we inverted the core mapping rule such that higher hydrophobicity was mapped to higher musical pitches (note = round(base note + (*H × S*))), while all other rules remained identical.
2. **Semantic Ablation Model:** To isolate the effect of the 2D representation structure itself, we created a model with a fixed, but random, mapping between the 20 amino acids and a set of 20 distinct pitches. This removes any biophysical meaning from the pitch assignment while preserving a consistent 1D-to-2D transformation.

### Biophysical Validation of the Sequence-to-Dynamics Mapping Principle

To establish a rigorous biophysical foundation for our entire sonification framework, we performed a critical computational experiment. The objective was to validate our core premise: that it is valid and meaningful to map static sequence information onto a dynamic, temporal medium like music. We hypothesized that if our encoding philosophy is sound, then the static features we use as proxies for dynamics (like B-factors for flexibility) must correlate with “ground truth” physical dynamics observed in simulation.

### Molecular Dynamics Simulations

To assess the generalizability of our findings, we performed molecular dynamics (MD) simulations on four proteins representing highly diverse structural fold families: T4 Lysozyme (PDB ID: 2LZM, an *α/β* protein), a de novo designed TIM-barrel (PDB ID: 5BVL, an *α/β* protein), Villin Headpiece (PDB ID: 2F4K, an all-*α* protein), and Green Fluorescent Protein (PDB ID: 1EMA, an all-*β* barrel protein). All simulations were conducted using the GROMACS package with the AMBER99SB-ILDN force field and the TIP3P water model. Following standard system preparation and equilibration, a 100 ns production MD simulation was carried out for each system in the NPT ensemble. The stability of each trajectory was confirmed by the convergence of the backbone root-mean-square deviation (RMSD). A detailed simulation protocol is provided in the Supplementary Information.

### Data Analysis

For each trajectory, the Root-Mean-Square Fluctuation (RMSF) of each residue’s C*α* atom was calculated from the full 100 ns production run using the `gmx rmsf` tool, serving as the ground-truth measure of dynamic flexibility. The experimental B-factors were extracted from the corresponding PDB files. Given that these two metrics are theoretically related [54], we quantified their linear relationship by calculating the Pearson correlation coefficient (r) and its corresponding p-value. This direct comparison validates the link between the static crystallographic data and dynamic behavior.

### Comparative Models for Protein Function Classification

Our analytical pipeline was designed to rigorously investigate the interplay between the nature of the input signal (1D vs. 2D) and the model’s learning paradigm. All sonification-based models were trained using data generated by the three mapping schemes described previously, with performance in comparative tables corresponding to the best-performing Theory-Driven model unless stated otherwise.

### Baseline Models on 1D Physicochemical Signals

We first established performance baselines using the primary 1D physicochemical sequence. Each protein sequence was converted into a numerical signal by mapping amino acids to their Kyte-Doolittle hydrophobicity values.

#### Classical ML with Spectral Features

We applied a Fast Fourier Transform (FFT) to each 1D signal to extract spectral features, which were then used to train an ensemble machine learning model.

#### 1D Convolutional Neural Network

We also applied a 1D Convolutional Neural Network (CNN) directly to the 1D signals to establish a modern deep learning baseline.

#### Sonification-Based Models on 2D Spectrograms

Next, we advanced the data representation by translating protein sequences into 2D spectrograms via sonification.

#### Classical ML with Acoustic Features

To isolate the impact of the 2D representation, we extracted Mel-Frequency Cepstral Coefficients (MFCCs) from each spectrogram and fed them into the same classical ML pipeline.

#### Model Architecture and Ablation Study

Finally, we combined the advanced 2D representation with a sophisticated, end-to-end deep learning model. The final model features a dual-branch architecture, fusing features from a **Visual Branch** (a ConvNeXt-Tiny backbone processing the spectrogram image [55]) and an **Acoustic Branch** (a GRU with attention processing the MFCC sequence). The concatenated features are passed to a final classification head.

To deconstruct the contributions of each modality and investigate the source of the model’s performance, we designed two additional model configurations for an ablation study. The **Visual-Only Model** consists solely of the ConvNeXt-Tiny backbone, processing the complete spectrogram as an image. Conversely, the **Acoustic-Only Model** utilizes only the GRU with attention mechanism, taking the MFCC sequence derived from the spectrogram as its input. These configurations allow for a direct comparison of the predictive power inherent in the visual versus the acoustic interpretation of the sonified data.

### Benchmark Models for Comparison

To situate our model’s performance, we benchmarked it against prominent protein language models (PLMs) and classical homology-based methods.

#### Protein Language Models (ESM-2 and ProtBERT)

We evaluated two prominent PLMs: ESM-2 (8M [56] and 35M [57] parameter versions) [5] and ProtBERT [6]. Both were tested under two conditions: a **fine-tuning** setting using official pre-trained weights, and a **from-scratch** training setting using only our dataset, which provides a direct comparison of data efficiency and the inherent capabilities of the architectures.

#### Homology-Based Search (BLAST+kNN)

As a classical baseline, we implemented two BLAST-based k-Nearest Neighbors (kNN) classifiers [58, 59]. The primary benchmark searches against the manually curated **Swiss-Prot database** to simulate a real-world annotation scenario. A secondary baseline searches against our own training set, serving as a measure of performance based on sequence similarity within our specific data distribution. We report the more challenging Swiss-Prot results in the main text.

### Generative Design of GFP Variants

To demonstrate the practical utility of our encoding, we integrated it into a conditional diffusion model to guide the *de novo* design of Green Fluorescent Protein (GFP) variants. The model was trained on experimental fluorescence data [60] and employed a multi-objective selection strategy that balanced predicted fitness with a harmonic score derived from our sonification framework, aiming to generate novel sequences that were both high-functioning and structurally viable.

## Results

### Biophysical Foundation of the Sonification Framework

Before evaluating classification models, we first sought to validate the core premise of our quantitative sonification framework: that mapping static sequence properties to a dynamic medium is a biophysically meaningful approach. To address the potential limitations of a single-protein validation, we extended our analysis to four proteins representing diverse fold families (all-*α*, all-*β*, and two distinct *α/β* topologies), each simulated for a comprehensive 100 ns.

Our analysis revealed statistically significant positive correlations between the experimental B-factors and the simulation-derived RMSF values for all four proteins, as detailed in Table 1. Notably, the all-*α* Villin Headpiece (2F4K) showed a strong correlation (r = 0.9030), while even the more complex proteins like the TIM-barrel (5BVL) and T4 Lysozyme (2LZM) demonstrated significant correlations. These results provide quantitative evidence supporting the use of static crystallographic data as a proxy for dynamic flexibility across different protein architectures. This supports the premise that the fundamental principle of our encoding is grounded in the physical reality of protein dynamics, strengthening the biophysical rationale for our methodology. The correspondence is visualized in Figure 2.

**Table 1:** Correlation between experimental B-factors and MD-derived RMSF across diverse protein folds.

| PDB ID | Protein Fold Family | Pearson’s r | P-value |
| --- | --- | --- | --- |
| 2F4K | All-α (Villin Headpiece) | 0.9030 | 6.58e-13 |
| 5BVL | $\alpha/\beta$ (TIM-Barrel) | 0.6768 | 1.85e-25 |
| 2LZM | $\alpha/\beta$ (T4 Lysozyme) | 0.5478 | 3.15e-14 |
| 1EMA | All-β (GFP) | 0.3653 | 4.01e-08 |

**Figure 1.**
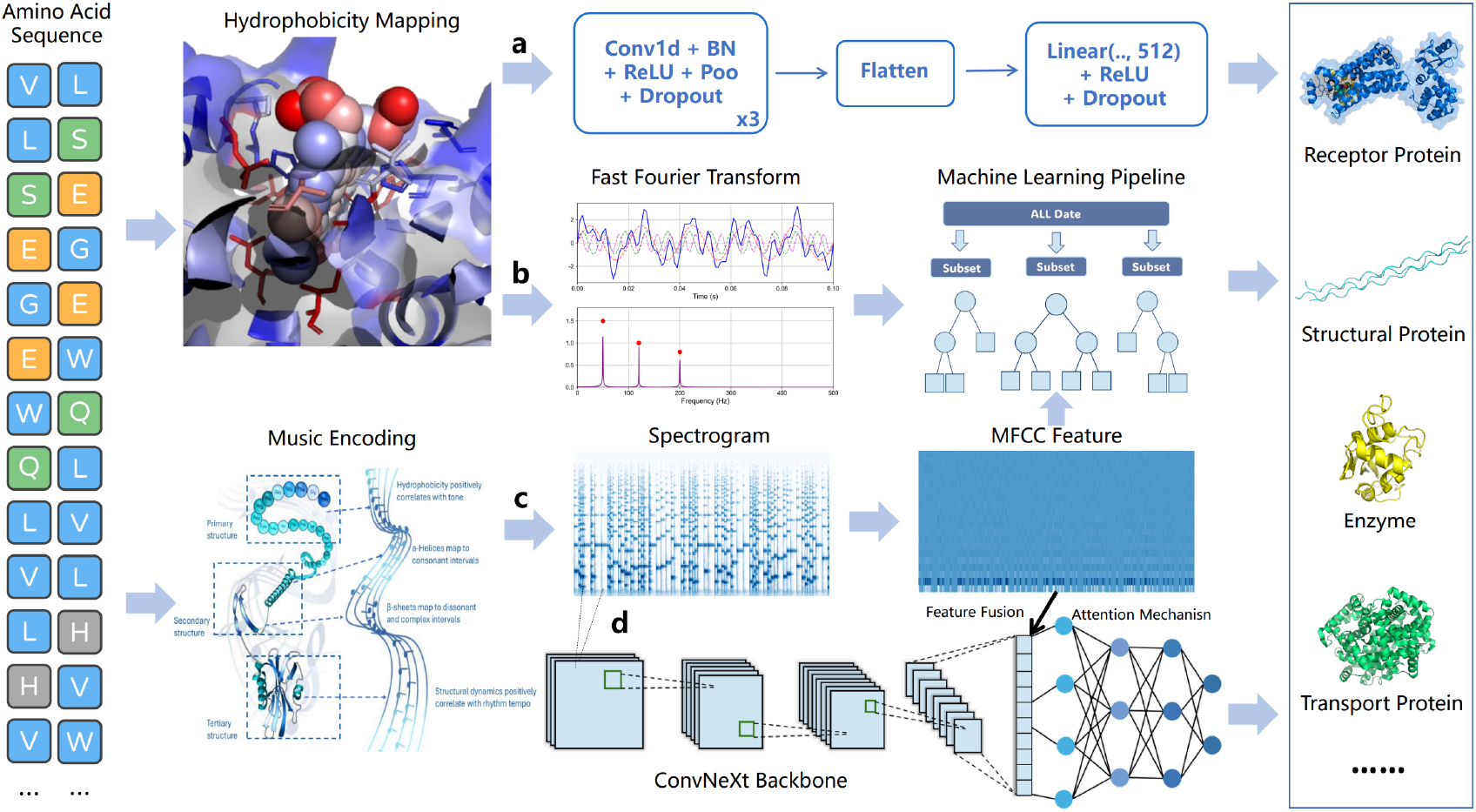
Our analytical pipeline is designed as a systematic comparative study to demonstrate the progressive benefits of evolving both data representation and model complexity. The workflow encompasses four distinct models: (a) 1D CNN on 1D Signal: A deep learning model that learns features directly from the raw 1D physicochemical signal. (b) Classical ML on 1D Signal: A baseline model that relies on spectral features extracted via Fast Fourier Transform (FFT) from the 1D signal. (c) Classical ML on 2D Spectrogram: A model that uses engineered audio features (Melfrequency cepstral coefficients, MFCC) from spectrograms. (d) End-to-End Fusion Model on 2D Spectrogram: Our final model, which employs a Convolutional Neural Network (CNN) and an attention mechanism to automatically learn and fuse features from the 2D spectrograms.

**Figure 2.**
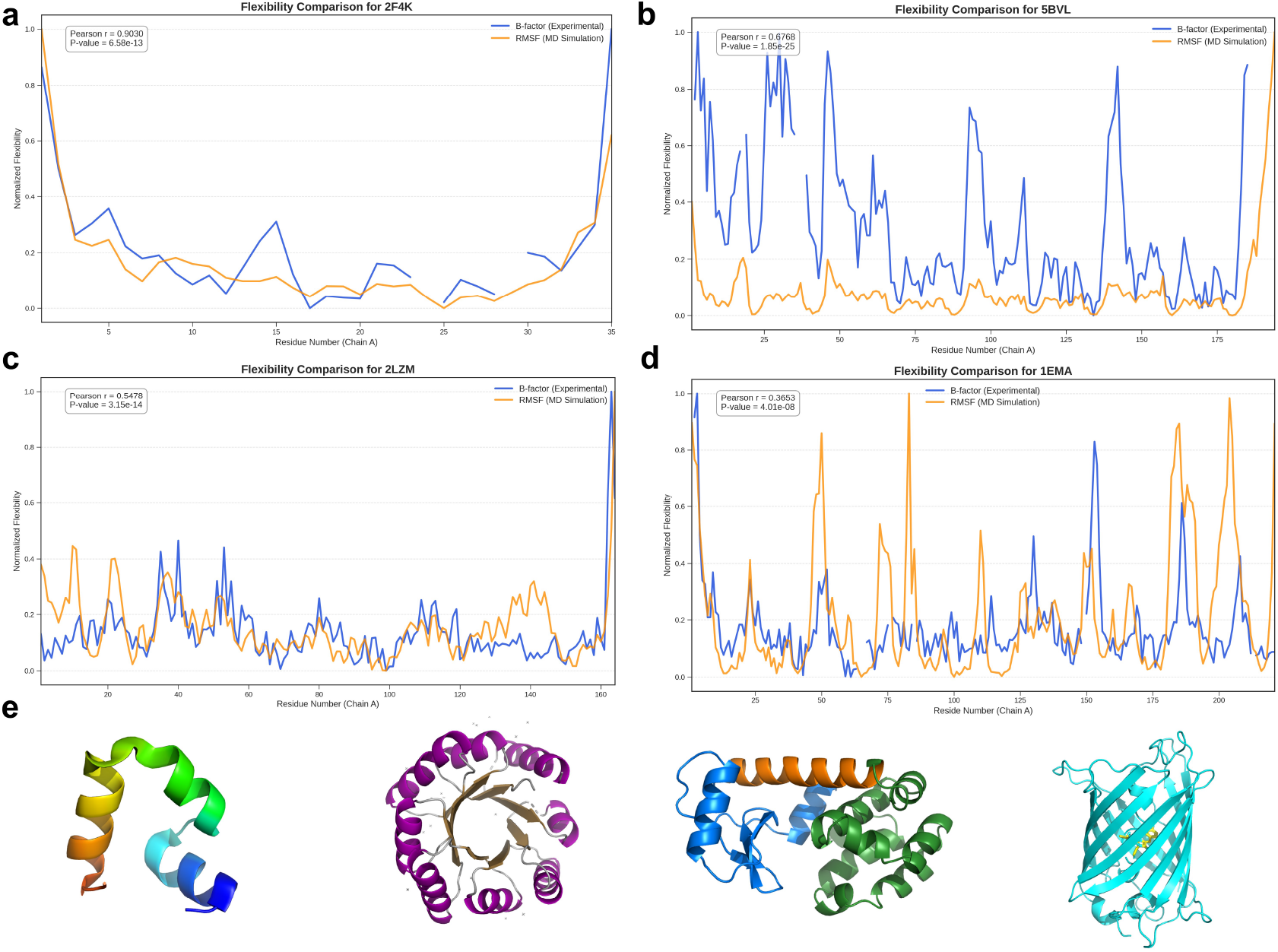
Assessing B-factors as a Proxy for Dynamic Flexibility Across Diverse Protein Fold Families. Comparison of experimental flexibility (crystallographic B-factors, blue) and simulated dynamic flexibility (RMSF from 100 ns MD simulations, orange) for four proteins representing distinct structural classes. The plots show statistically significant positive correlations for (a) the all-*α* Villin Headpiece (2F4K, r=0.903), (b) the *α/β* TIM-Barrel (5BVL, r=0.677), (c) the *α/β* T4 Lysozyme (2LZM, r=0.548), and (d) the all-*β* GFP (1EMA, r=0.365). All correlations are highly significant (p *<* 1e-7). This analysis provides biophysical support for the principle of mapping static structural properties to dynamic features in our sonification framework. (e) Cartoon representations of the protein structures analyzed in (A-D), shown in corresponding order from left to right.

### Visual and Quantitative Interpretation of the Protein Score

To provide an intuitive illustration of our encoding framework, we translated protein sequences not only into spectrograms for machine analysis but also into conventional musical scores for human interpretation. This allows for a direct visual inspection of how a protein’s physicochemical properties are mapped onto musical language. To illustrate this, we selected two proteins with contrasting architectures: the small, highly stable, all-alpha-helical Villin Headpiece (1VII), and the larger, structurally complex T4 Lysozyme (2LZM), which contains a mix of helices, sheets, and flexible loops.

As shown in Figure 3, the resulting scores are visually and quantitatively distinct, reflecting their underlying biochemical differences. The score for the stable Villin Headpiece (Figure 3a) exhibits a higher degree of regularity. Its melody, determined by amino acid hydrophobicity, shows less dramatic fluctuation, and its rhythm, governed by molecular weight, is more uniform. In contrast, the score for the complex T4 Lysozyme (Figure 3b) is noticeably more varied, featuring greater melodic leaps and more intricate rhythmic patterns.

**Figure 3.**
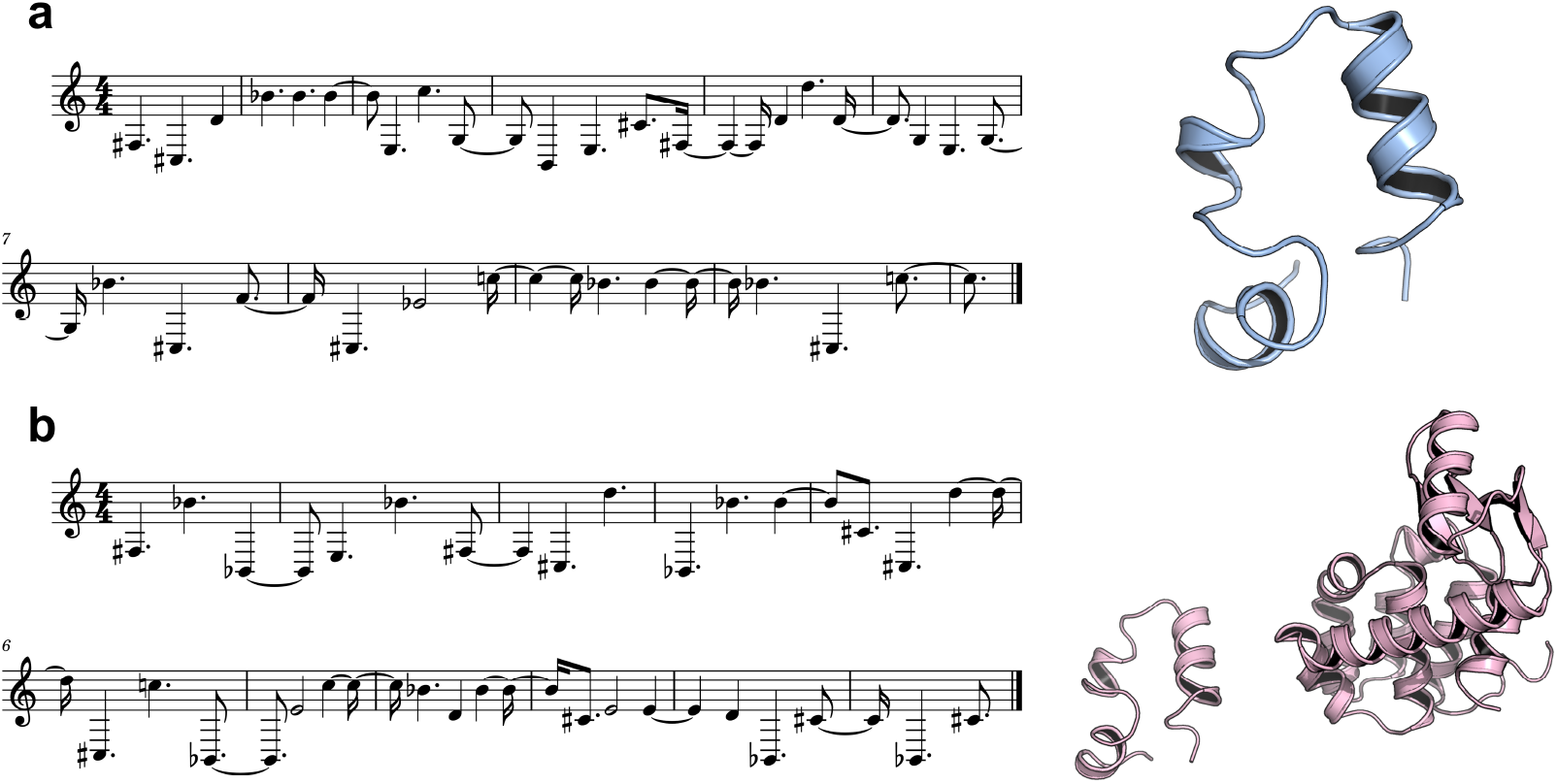
Visualization of Protein-to-Music Translation as Sheet Music. Musical scores were generated from the first 30 amino acids of two representative proteins. (a) Villin Headpiece (1VII): This structurally stable protein translates into a score with a constrained melodic range and regular rhythm. (b) T4 Lysozyme (2LZM): This complex protein produces a score with greater melodic leaps and more intricate rhythms. These visual differences are quantitatively supported: the T4 Lysozyme fragment exhibits a 35.1% higher melodic complexity score and a 7.4% higher rhythmic complexity score compared to the Villin fragment. This is consistent with the greater diversity in its physicochemical properties.

To formalize this visual observation, we calculated complexity metrics for the first 30-amino-acid fragment. The ‘melodic complexity’ was quantified as the average absolute difference in hydrophobicity between adjacent residues, while ‘rhythmic complexity’ was defined as the standard deviation of molecular weights. The analysis indicates that the T4 Lysozyme fragment is more complex, possessing a 35.1% higher melodic complexity score (4.58 vs. 3.39) and a 7.4% higher rhythmic complexity score (29.70 vs. 27.65) than the Villin fragment. This visual and quantitative comparison suggests that our framework can translate core structural and chemical features into distinct, interpretable musical patterns.

### Beyond Semantics: The Role of Representational Structure

To ascertain whether the predictive power of sonification stems from the specific semantic encoding rules or the underlying structural transformation, our investigation was conducted on the comprehensive 12-class, 18,000-sequence dataset. The results, summarized in Table 2, provide quantitative evidence suggesting that the structural transformation from a 1D sequence to a 2D spectrogram is a major contributor to the observed performance improvement.

Our theory-driven quantitative model, which incorporates biophysical information, achieved an accuracy of 84.00%. However, the most striking finding is that the control models with inverted and random semantics still achieved remarkably high accuracies of 82.69% and 81.08%, respectively. All three 2D sonification-based models substantially outperformed our strongest 1D baseline (61.31%, see Table 3). This result indicates that the vast majority of the predictive power is unlocked by the 2D representation itself, which enables the deep learning model to learn long-range dependencies and complex textural patterns. While the structural transformation is primary, biophysical semantics provide a consistent and measurable performance advantage (a nearly 3-point gain from the random to the theory-driven model), suggesting they offer a valuable inductive bias rather than being an absolute requirement.

**Table 2:** Performance comparison of different sonification mapping schemes on the 12-class dataset. All models use the identical end-to-end fusion architecture and were evaluated on the same independent test set to deconstruct the source of performance.

| Mapping Scheme | Core Principle | Accuracy (%) | Precision | Recall | F1-Score |
| --- | --- | --- | --- | --- | --- |
| **Theory-Driven** | Biophysical Semantics | **84.00** | **0.84** | **0.84** | **0.84** |
| Inverted Semantics | Inverted Biophysics | 82.69 | 0.83 | 0.83 | 0.83 |
| Semantic Ablation (Random) | Structure-Only | 81.08 | 0.81 | 0.81 | 0.81 |

**Table 3:** Comparative Performance of Classification Models on the Independent Test Set, demonstrating the impact of evolving data representation (1D vs. 2D) and model paradigm.

| Data | Model | Accuracy (%) | Precision | Recall | F1-Score |
| --- | --- | --- | --- | --- | --- |
| **1D Signal** | FFT + Classical ML | 45.86 | 0.45 | 0.46 | 0.43 |
|  | **1D CNN (Deep Learning)** | **61.31** | **0.61** | **0.61** | **0.60** |
| **2D Spectrogram** | MFCC + Classical ML | 69.47 | 0.69 | 0.69 | 0.69 |
|  | **Fusion Model (Deep Learning)** | **84.00** | **0.84** | **0.84** | **0.84** |

### The Synergy of Representation and Architecture

Building on the finding that 2D sonified representations are effective, we conducted a broader comparative analysis to understand the synergy between data representation and model architecture. The results, summarized in Table 3, reveal a clear trend on our 12-class benchmark.

Our investigation began with the 1D physicochemical signal. The classical approach (FFT + ML) achieved an accuracy of 45.86%. Applying a modern 1D CNN improved performance to 61.31%. We then advanced the data representation to 2D spectrograms using our theory-driven mapping. A classical ML pipeline with engineered MFCC features achieved 69.47% accuracy, a significant improvement over both 1D methods, confirming the intrinsic value of the 2D representation. The highest performance was achieved by pairing the advanced representation with an advanced model: our end-to-end fusion model reached the final accuracy of 84.00%. This progression suggests that high performance is unlocked when an enriched 2D representation is paired with a deep learning architecture capable of exploiting its complexity.

### Ablation Study: Deconstructing the Source of Predictive Power

To further investigate the individual contributions of the visual and acoustic feature streams, we conducted an ablation study. As shown in Table 4, our full fusion model achieved the highest accuracy at 84.00%. Notably, the single-branch models also demonstrated substantial predictive capability. The Visual-Only model, relying on the ConvNeXt backbone to process the raw spectrogram, achieved 76.56% accuracy, while the Acoustic-Only model, using the GRU to process extracted MFCC features, reached 77.17% accuracy.

The strong performance of both unimodal configurations suggests that the 2D spectrogram generated through sonification is an information-rich representation. It indicates that functionally relevant patterns are encoded in a manner that is accessible to both direct visual analysis (as an image) and acoustic feature extraction (as a temporal signal). This finding implies that the effectiveness of our approach is not solely dependent on the final fusion architecture, but is fundamentally rooted in the descriptive power of the 2D representation itself.

**Table 4:**
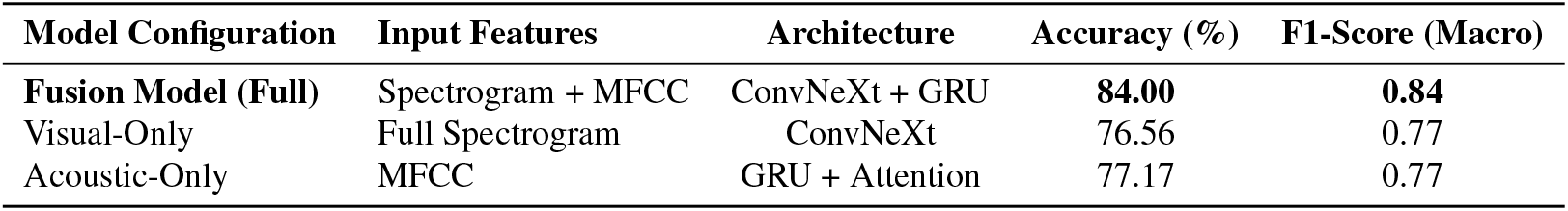
Ablation study of the fusion model on the independent test set. Performance of the full model is compared against its individual visual and acoustic branches.

| Model Configuration | Input Features | Architecture | Accuracy (%) | F1-Score (Macro) |
| --- | --- | --- | --- | --- |
| **Fusion Model (Full)** | Spectrogram + MFCC | ConvNeXt + GRU | **84.00** | **0.84** |
| Visual-Only | Full Spectrogram | ConvNeXt | 76.56 | 0.77 |
| Acoustic-Only | MFCC | GRU + Attention | 77.17 | 0.77 |

### Comparative Benchmarking Against Established Methods

To situate our model’s performance in a broader context, we benchmarked our final fusion model against established protein language models (ProtBERT and ESM-2) and homology-based search methods. The results are detailed in Table 5 and visualized in Figure 4.

**Table 5:**
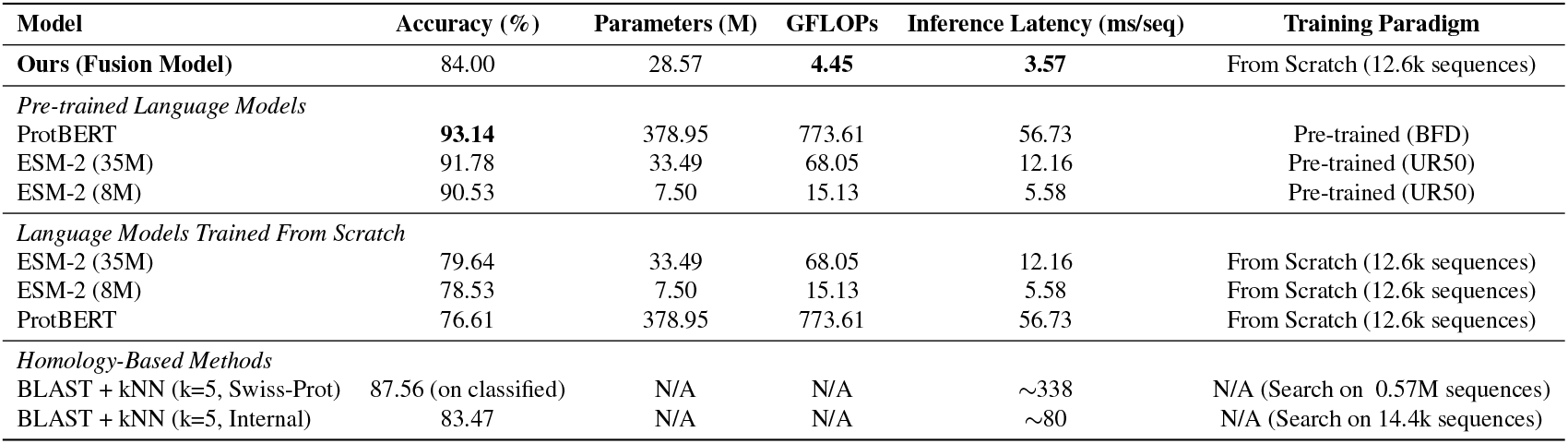
Comparison of performance and computational complexity against state-of-the-art models.

**Figure 4.**
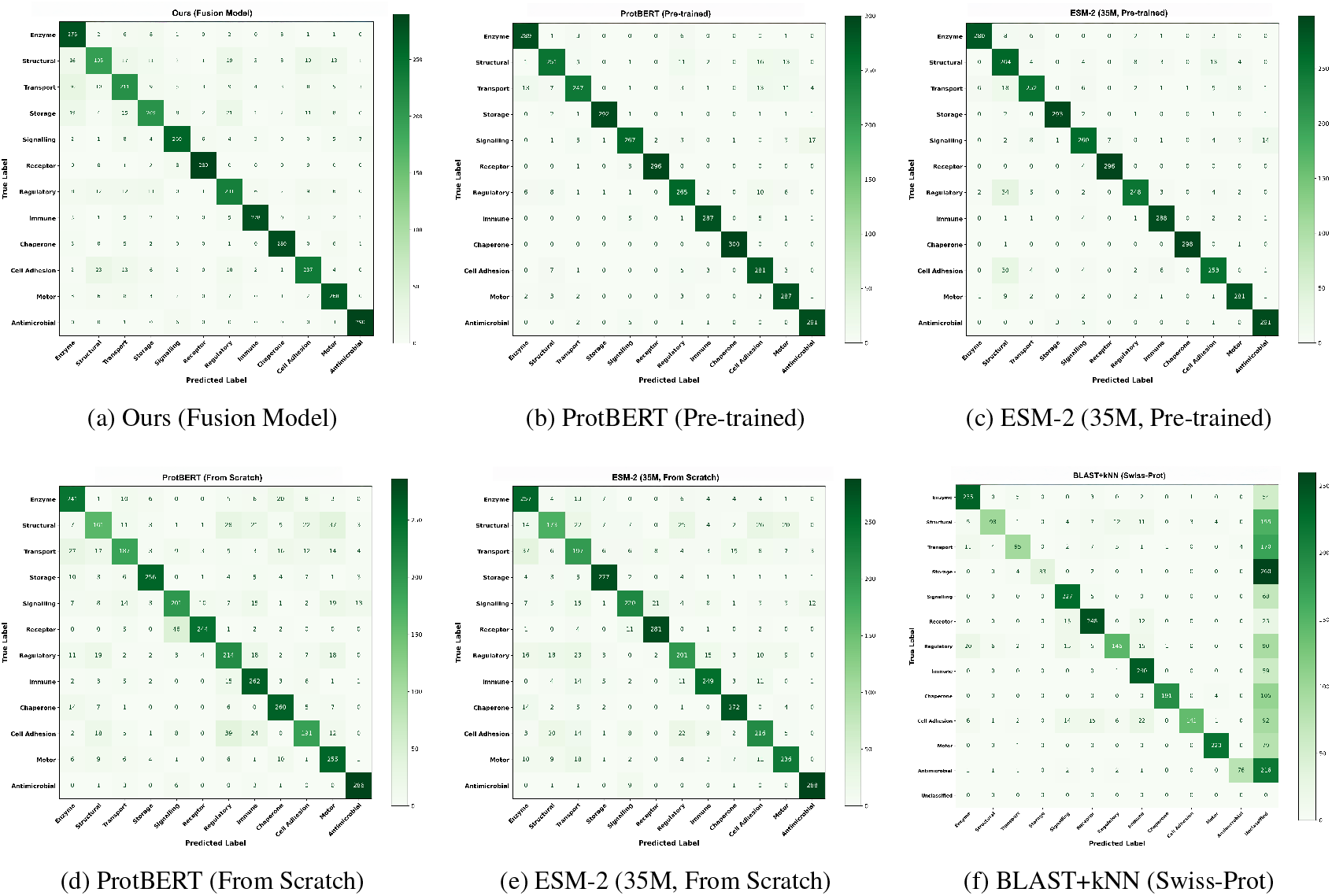
Comparative Confusion Matrices on the independent 12-class test set. The top row compares our model (a) against high-performing pre-trained models: ProtBERT (b) and ESM-2 (c). The bottom row directly compares models trained from scratch—ProtBERT (d) and ESM-2 (e)—and the homology-based baseline (f), highlighting our model’s data efficiency.

As expected, large pre-trained models achieved the highest overall accuracies, with ProtBERT reaching 93.14% and ESM-2 reaching up to 91.78%. This outcome is attributable to their extensive pre-training on massive external datasets (e.g., BFD [61] and UR50 [62]), which provides them with extensive prior knowledge derived from evolutionary and structural patterns.

However, a more direct comparison of model architecture and representational efficiency emerges when this pretraining advantage is removed. When trained from scratch on our 12,600-sequence training set, our fusion model (84.00%) achieved a higher accuracy than the transformer-based architectures tested. It surpassed the ESM-2 models (78.53%-79.64%) and, notably, also the much larger ProtBERT model (76.61%). This finding suggests that, for the task and dataset presented here, our sonification process may offer a data-efficient representation, allowing a vision-based model to learn relevant patterns effectively under data-limited conditions. This indicates that under these data-limited conditions, a vision-based model using our spectrogram representation was able to learn relevant functional patterns more effectively than the tested transformer architectures could from raw 1D sequences.

The homology-based BLAST+kNN approach against the Swiss-Prot database achieved a high accuracy of 87.56%, however, this was only on the 61.86% of test sequences for which it could find reliable homologs, leaving the remaining 38% unclassified. As an internal control, applying the same method against our own training set yielded an accuracy of 83.47% across all test sequences. This comparison indicates that while our model holds a slight performance edge over the internal BLAST, a significant portion of the classification task within this dataset can be solved by sequence similarity alone. Thus, our model offers a viable alternative that does not rely on finding close homologs in a reference database, suggesting its potential utility for sequences that lack clear evolutionary relatives.

Latency for our model and PLMs was measured as the average of 100 runs on a single NVIDIA 4090 24GB GPU. The BLAST+kNN latencies were measured on a system with a 20 vCPU Intel® Xeon® Platinum 8470Q processor and are reported per query sequence. GFLOPs for PLMs are calculated for a sequence length of 1024. “From Scratch” models were trained on our dataset (12,600 sequences) without any pre-training. Pre-trained models utilize weights trained on large external datasets like BFD or UR50. BLAST accuracy for the Swiss-Prot search is reported only on the 61.86% of test sequences for which homologs could be found.

### Validation on an External Enzyme Classification Benchmark

To evaluate the generalizability of our framework, we tested our approach on a larger dataset constructed from the external CARE enzyme classification benchmark [49]. For this validation, we retrained our fusion model from scratch on a new dataset of 35,000 sequences. The model, still employing our theory-driven mapping, achieved a competitive overall accuracy of 90.44% on the 7,000-sequence test set (Table 6). This strong performance on a large-scale, independent benchmark demonstrates that our sonification-based approach is not overfitted to our initial dataset and generalizes well to new classification tasks.

**Table 6:** Performance of the Fusion Model on the CARE Benchmark Test Set.

| Metric | Precision | Recall | F1-Score | Accuracy (%) |
| --- | --- | --- | --- | --- |
| Macro Average | 0.91 | 0.90 | 0.90 | 90.44 |
| Weighted Average | 0.91 | 0.90 | 0.90 |  |

### Generative Validation: De Novo Design of Viable GFP Variants

To demonstrate the practical utility of our encoding in a generative context, we integrated our framework into a conditional diffusion model to guide the *de novo* design of Green Fluorescent Protein (GFP) variants. Our goal was to balance the dual objectives of high functional fitness and sequence diversity, as selecting solely for the highest fitness can lead to a narrow set of solutions. To achieve this, we implemented a two-stage selection strategy. First, we identified an elite pool of candidates with the highest predicted fitness, ensuring a baseline of high performance. Then, within this elite pool, we selected the final variants using a composite score that weighed both predicted fitness and the harmonic score derived from our sonification framework.

As shown in Figure 5, this strategy yielded promising results. Compared to a fitness-only selection method, our approach generated a population of variants that not only maintained high fitness levels but also exhibited greater sequence diversity and explored a broader region of the harmonic score landscape. To validate the structural integrity of these designs, computational analysis using ESMFold was performed. The results (Figure 6) indicated that the generated variants largely retained the canonical GFP *β*-barrel fold with high confidence and low RMSD values (0.062–0.265 Å) relative to the wild-type, providing evidence that our sonification-derived harmonic score may serve as a useful proxy for structural viability in this context.

**Figure 5.**
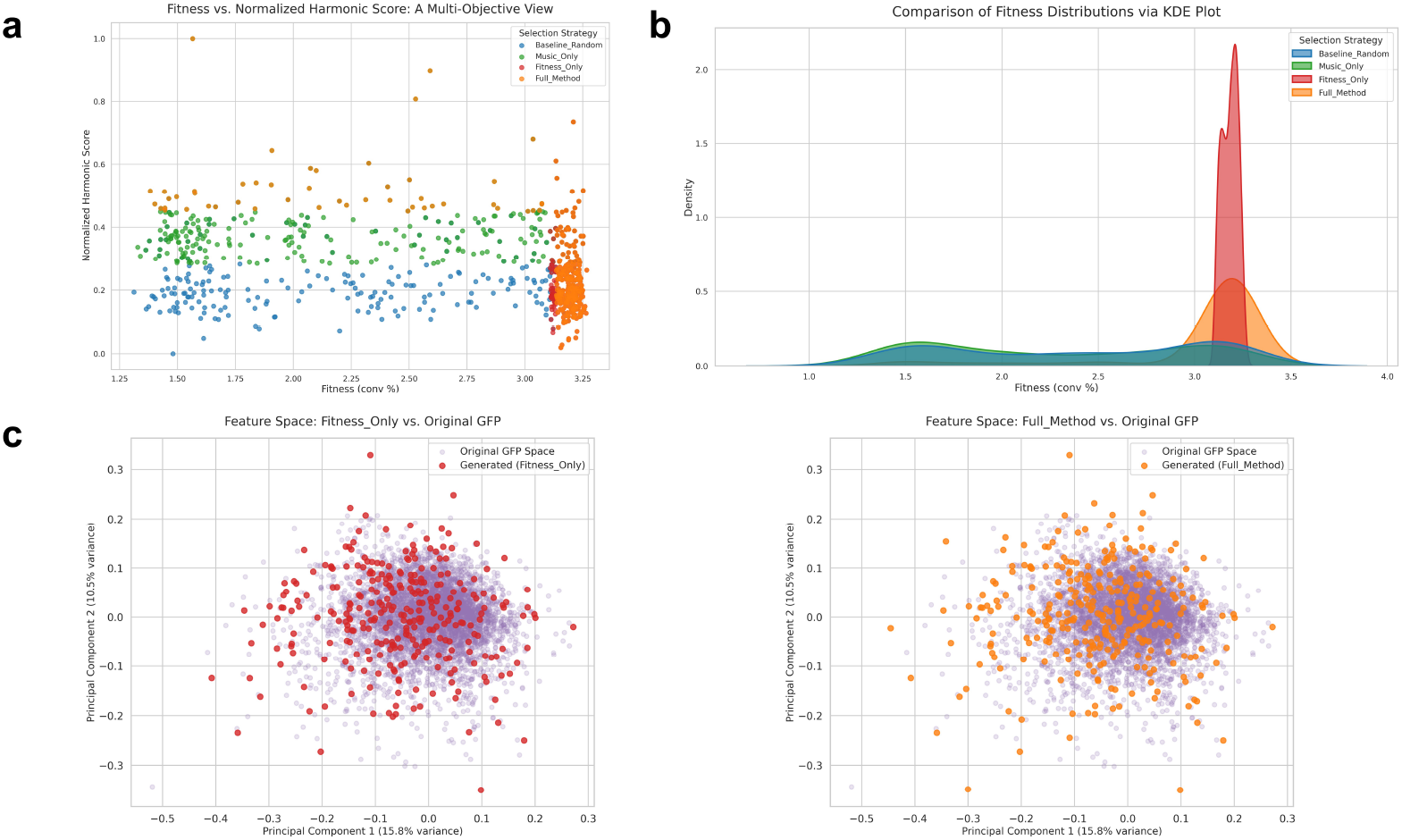
Comparison of Selection Strategies for Multi-Objective Protein Optimization (A) Multi-Objective Land-scape: This scatter plot shows the relationship between predicted fitness and normalized harmonic score. The `Full_Method` (orange) explores regions of higher harmonic scores, achieving a mean harmonic score of 0.8396 *±* 0.0029. In contrast, the `Fitness_Only` population (red) is concentrated in a zone of high fitness (mean = 3.1831 *±* 0.0404) but is constrained to a narrow range of lower harmonic values (mean = 0.8386 *±* 0.0018). (B) Fitness Distributions: The Kernel Density Estimate (KDE) plot compares the fitness distributions. The `Full_Method` exhibits a significantly broader distribution (fitness = 3.0271 *±* 0.4414) compared to the sharply peaked `Fitness_Only` population (fitness = 3.1831 *±* 0.0404). The large standard deviation of the `Full_Method` quantitatively reflects the mixed nature of its population, comprising both high-performance variants and other sequences that contribute to its exploratory capability. (C) Sequence Space Visualization: Principal Component Analysis (PCA) visualizes the distribution of selected sequences. The `Fitness_Only` population forms a compact cluster, corresponding to a diversity score of 7.0261%. The `Full_Method` population occupies a visibly larger area of the sequence space, achieving a higher diversity score of 7.1159% and suggesting its ability to generate a more varied set of high-performing sequences.

**Figure 6.**
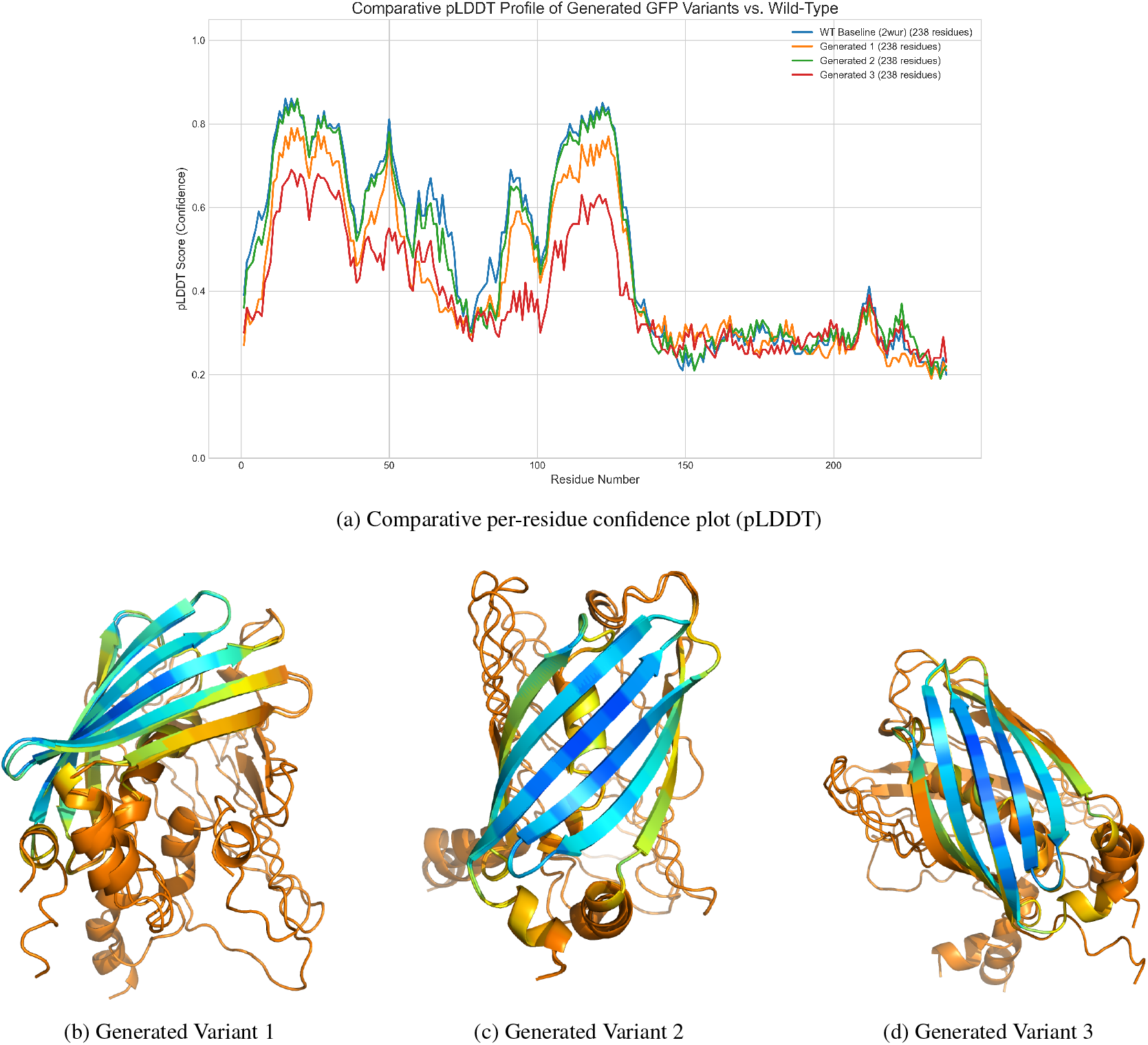
Computational validation of generated GFP variants. (a) Per-residue confidence scores (pLDDT) from ESMFold for the wild-type baseline and three generated variants. (b-d) Predicted 3D structures of the three variants, colored by pLDDT score (Blue: *>*0.9, Orange: *<*0.5). The generated variants appear to retain the essential *β*-barrel fold.

## Discussion

A central question motivating this study was whether our encoding rules were arbitrary or essential for performance. Our results provide a nuanced answer: the primary benefit of sonification appears to stem from the structural transformation of a 1D sequence into a 2D, information-dense representation. The strong performance of the semantic ablation model (81.08% accuracy) suggests that a deep learning model can learn functional patterns from the spectrogram’s structure alone, treating them as emergent properties. This ability to extract meaningful regularities from a complex, semantically-arbitrary representation resonates with neurophysiological findings on pattern recognition in uncertain musical contexts [63]. However, this does not render the encoding rules meaningless. The incremental performance gains from the inverted (82.69%) and our theory-driven (84.00%) models indicate that biophysically-grounded semantics provide a valuable inductive bias, an idea supported by our molecular dynamics simulations which validate the premise that mapping static sequence properties to a dynamic medium is a physically meaningful approach. This reframes our method within a broader intellectual tradition of cross-pollination between biology and music, where such analogies have been used both for analysis and the creation of novel bio-inspired materials [64–66].

This 1D-to-2D transformation is a powerful and established principle, not a new hypothesis. Its success is a cornerstone of landmark models like AlphaFold, which converts sequence information into 2D distance maps to capture the spatial relationships between all residue pairs [14, 15]. The same paradigm is fundamental to state-of-the-art methods in genomics [16, 17] and RNA secondary structure prediction [18], where predicting 2D interaction maps from 1D sequences is crucial for modeling long-range interactions (LRIs). This principle is not confined to biology; it is a core strategy in machine learning for capturing LRIs, for instance in graph neural networks (GNNs) where multi-scale representations have yielded order-of-magnitude performance gains on LRI-centric benchmarks [67, 68]. By “folding” a sequence into a spectrogram, we bring distant residues into proximity, allowing a standard CNN to model these global dependencies efficiently, without the quadratic cost of attention mechanisms [69]. This echoes other computational paradigms where function is derived from global network properties [70].

The richness of this representation is further evidenced by our ablation study. The strong standalone performance of both the Visual-Only and Acoustic-Only models demonstrates that functional information is encoded in a way that is accessible through different analytical lenses. This aligns with a growing body of work showing sonification can be a valuable complement to purely visual methods for data discovery in the life sciences [71, 72], and is conceptually analogous to how the mammalian auditory cortex utilizes joint spectro-temporal features for robust sound recognition [73].

Positioning our work relative to state-of-the-art Protein Language Models (PLMs) highlights a trade-off. While large, pre-trained models like ProtBERT and ESM-2 achieve superior accuracy, their performance advantage is removed when training is restricted to our dataset. Our fusion model’s competitive performance when trained from scratch suggests that the 2D representation is highly data-efficient for this task. This efficiency, however, comes with the overhead of a two-step process involving pre-conversion of sequences to spectrograms. This points toward a promising future direction: a truly end-to-end architecture, such as a Protein Spectrogram Transformer (PST). Inspired by the audio spectrogram transformer [74], a PST could learn to generate an optimal 2D representation directly from the 1D sequence, combining the representational power of our approach with the end-to-end learning paradigm of PLMs.

We acknowledge the limitations of our study. Our dataset, while functionally diverse, is modest by the standards of large-scale pre-training—a well-recognized challenge often necessitating customized approaches or domain adaptation techniques [75]. Similarly, our generative experiments on GFP, while a compelling proof-of-concept, require validation across a broader range of protein families to establish the generalizability of our harmonic score as a proxy for structural viability. Nevertheless, this initial validation is critical, as it suggests that features derived from our framework could serve as a practical regularizer in the *de novo* protein design cycle, where ensuring a viable three-dimensional fold is a cornerstone of success [76, 77].

In summary, this work establishes protein sonification as a viable strategy for creating computationally efficient and information-dense representations for function prediction. The primary benefit stems from the structural shift to a 2D format, which allows deep learning models to learn emergent functional patterns, with biophysical semantics providing a beneficial, but secondary, guiding principle.

## Conclusion

In this study, we developed and evaluated a quantitative sonification framework as a novel method for protein analysis. Our work demonstrates that translating protein sequences into 2D spectrograms creates a feature-rich representation that is effective for function prediction, showing signs of data efficiency and generalizability across different protein classification tasks. Through a series of systematic experiments, we provide evidence that the primary source of the model’s predictive power stems from the structural transformation of 1D sequence data into a 2D format. This allows for the capture of complex patterns as emergent properties, a process that is further enhanced, but not solely dependent on, biophysically-informed encoding rules. The utility of this representation extends beyond predictive tasks, as shown by our proof-of-concept where the encoding guided a generative model in designing novel, structurally viable GFP variants. This study contributes a systematic evaluation of protein sonification as a principled feature engineering strategy, highlighting its potential for both predictive and generative protein modeling. Ultimately, our work reinforces the paradigm of translating biological sequences into other domains, not merely for artistic inspiration [78], but as a structured approach for creating powerful, bio-inspired analytical methods and materials [65, 66].

## Data and Software Availability

The code is available at https://github.com/wyqmath/Symphony_of_Fate.

## Supporting Information

Supporting Information includes: a summary of data sources (Section A); detailed methodology of the protein-to-music sonification framework, including quantitative mapping rules, technical implementation for audio synthesis, and a discussion of alternative schemes (Section B); protocols for molecular dynamics (MD) simulations and RMSF analysis (Section C); complete implementation details for all classification systems, including the end-to-end fusion model and all benchmark models (1D-CNN, PLMs, homology-based) (Section D); implementation of the generative framework for GFP variants, detailing the fitness predictor, Harmonic Score, and directed evolution strategy (Section E); a case study on spectral feature analysis of protein structures (Section F); and all supporting figures and tables (Figures S1–S7; Tables S1–S6) (PDF).

## Author Contributions

**Yiquan Wang:** Conceptualization, Methodology, Software, Investigation, Writing – Original Draft, Writing – Review & Editing. **Minnuo Cai:** Software, Investigation, Visualization, Writing – Original Draft, Writing – Review & Editing. **Yuhua Dong:** Conceptualization, Methodology, Software, Investigation, Writing – Original Draft, Writing – Review & Editing. **Yahui Ma:** Formal Analysis, Writing – Original Draft. **Kai Wei:** Project Administration, Writing – Review & Editing.

## Declaration of Interests

The authors declare no competing interests

## Funding

This work was supported by the Natural Science Foundation of Xinjiang Uygur Autonomous Region (Grant Number: 2024D01C216) and the “Tianchi Talents” introduction plan.

## Acknowledgements

The authors extend their sincere gratitude to colleagues, mentors, and Shenzhen X-Institute for their invaluable discussions and support, and to the anonymous reviewers for their constructive feedback which significantly improved this manuscript. Sincere thanks are also extended to Ms. Chenyu Shen for her invaluable help with the musical scores. We also acknowledge the public databases that made this work possible, including the National Center for Biotechnology Information (NCBI), UniProt, BRENDA, TCDB, and APD.

## For Table of Contents Use Only

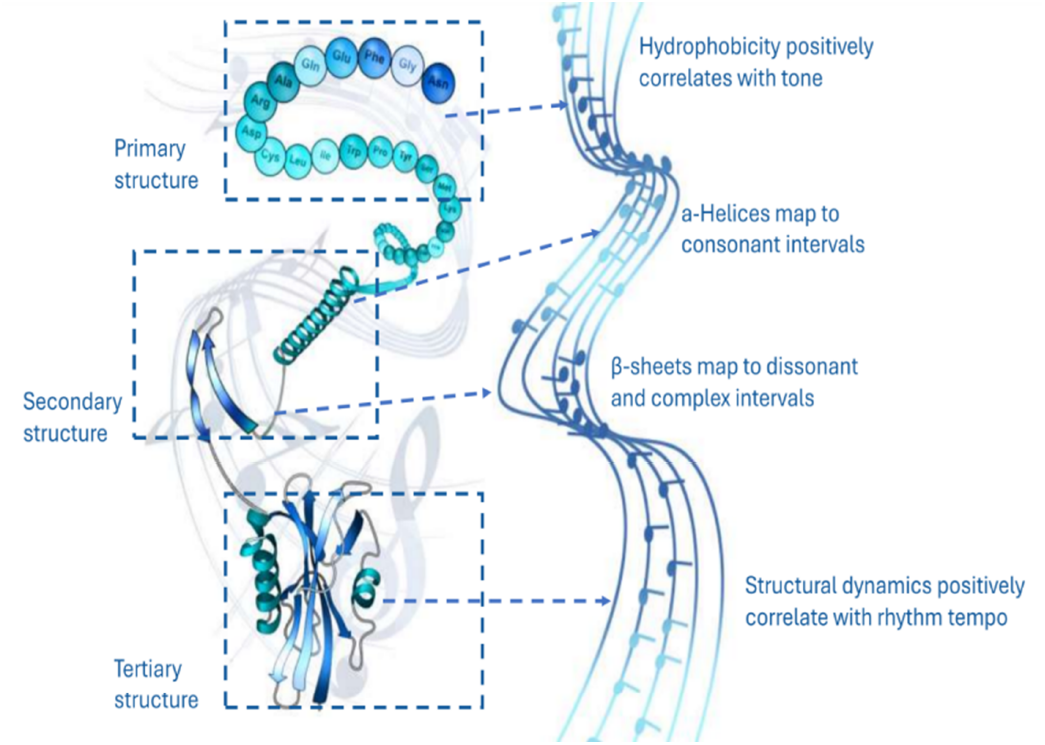

## Notes

### Competing Interest Statement

The authors have declared no competing interest.

## References

[1] H. Hegyi and M. Gerstein. The relationship between protein structure and function: a comprehensive survey with application to the yeast genome. Journal of molecular biology, 288(1): 147–164, 1999.

[2] M. I. Sadowski and D. T. Jones. The sequence–structure relationship and protein function prediction. Current opinion in structural biology, 19(3): 357–362, 2009.

[3] D. Whitford. Proteins: Structure and Function. John Wiley & Sons, 2013.

[4] J. Koehler Leman, P. Szczerbiak, P. D. Renfrew, V. Gligorijevic, D. Berenberg, T. Vatanen, T. Kosciolek, et al. Sequence-structure-function relationships in the microbial protein universe. Nature communications, 14(1): 2351, 2023.

[5] Zeming Lin, Halil Akin, Roshan Rao, Brian Hie, Zhongkai Zhu, Wenting Lu, Allan Smetanin, Robert Verkuil, Ori Kabeli, Yaniv Shmueli, Amir Ferdaos, Nevan Krogan, Tom Sercu, and Adam Rives. Evolutionary-scale prediction of atomic-level protein structure with a language model. Science, 379(6637): 1123–1130, 2023.

[6] Ahmed Elnaggar, Michael Heinzinger, Christian Dallago, Ghalia Rehawi, Yu Wang, Llion Jones, Tom Gibbs, Tamas Feher, Christoph Angerer, Martin Steinegger, Debsindhu Bhowmik, and Burkhard Rost. ProtTrans: Towards Cracking the Language of Life’s Code Through Self-Supervised Learning. IEEE Transactions on Pattern Analysis and Machine Intelligence, 44(10): 7112–7127, 2021.

[7] J. Jumper, R. Evans, A. Pritzel, T. Green, M. Figurnov, O. Ronneberger, and D. Hassabis. Highly accurate protein structure prediction with alphafold. Nature, 596(7873): 583–589, 2021.

[8] L. Wang, X. Li, H. Zhang, J. Wang, D. Jiang, Z. Xue, and Y. Wang. A comprehensive review of protein language models. arXiv preprint arXiv:2502.06881, 2025.

[9] T. Bepler and B. Berger. Learning the protein language: Evolution, structure, and function. Cell systems, 12(6): 654–669, 2021.

[10] J. Y. Chen, J. F. Wang, Y. Hu, X. H. Li, Y. R. Qian, and C. L. Song. Evaluating the advancements in protein language models for encoding strategies in protein function prediction: a comprehensive review. Frontiers in bioengineering and biotechnology, 13:1506508, 2025.

[11] C. H. Yu, Z. Qin, F. J. Martin-Martinez, and M. J. Buehler. A self-consistent sonification method to translate amino acid sequences into musical compositions and application in protein design using artificial intelligence. ACS Nano, 13(7): 7471–7482, 2019.

[12] M. J. Buehler. Unsupervised cross-domain translation via deep learning and adversarial attention neural networks and application to music-inspired protein designs. Patterns, 4(3), 2023.

[13] M. Milazzo, G. I. Anderson, and M. J. Buehler. Bioinspired translation of classical music into de novo protein structures using deep learning and molecular modeling. Bioinspiration & Biomimetics, 17(1): 015001, 2021.

[14] S. Wang, S. Sun, Z. Li, R. Zhang, and J. Xu. Accurate de novo prediction of protein contact map by ultra-deep learning model. PLoS computational biology, 13(1):e1005324, 2017.

[15] A. W. Senior, R. Evans, J. Jumper, J. Kirkpatrick, L. Sifre, T. Green, et al. Improved protein structure prediction using potentials from deep learning. Nature, 577(7792): 706–710, 2020.

[16] G. Fudenberg, D. R. Kelley, and K. S. Pollard. Predicting 3D genome folding from DNA sequence with Akita. Nature methods, 17(11): 1111–1117, 2020.

[17] R. Schwessinger, M. Gosden, D. Downes, R. C. Brown, A. M. Oudelaar, J. Telenius, …, and J. R. Hughes. DeepC: predicting 3D genome folding using megabase-scale transfer learning. Nature methods, 17(11):1118–1124, 2020.

[18] L. A. Bugnon, L. Di Persia, M. Gerard, J. Raad, S. Prochetto, E. Fenoy, …, and D. H. Milone. sincFold: end-to-end learning of short-and long-range interactions in RNA secondary structure. Briefings in Bioinformatics, 25(4), 2024.

[19] E. Nogales and S. H. Scheres. Cryo-em: A unique tool for the visualization of macromolecular complexity. Molecular Cell, 58(4): 677–689, 2015.

[20] K. M. Yip, N. Fischer, E. Paknia, A. Chari, and H. Stark. Atomic-resolution protein structure determination by cryo-em. Nature, 587(7832): 157–161, 2020.

[21] J. L. Klepeis, K. Lindorff-Larsen, R. O. Dror, and D. E. Shaw. Long-timescale molecular dynamics simulations of protein structure and function. Current Opinion in Structural Biology, 19(2): 120–127, 2009.

[22] M. AlQuraishi. Machine learning in protein structure prediction. Current Opinion in Chemical Biology, 65: 1–8, 2021.

[23] Y. Meng, Z. Zhang, C. Zhou, X. Tang, X. Hu, G. Tian, and Y. Yao. Protein structure prediction via deep learning: an in-depth review. Frontiers in Pharmacology, 16:1498662, 2025.

[24] M. Varadi, S. Anyango, M. Deshpande, S. Nair, C. Natassia, G. Yordanova, and S. Velankar. Alphafold protein structure database: Massively expanding the structural coverage of protein-sequence space with high-accuracy models. Nucleic Acids Research, 50(D1):D439–D444, 2022.

[25] S. Hertig, N. R. Latorraca, and R. O. Dror. Revealing atomic-level mechanisms of protein allostery with molecular dynamics simulations. PLoS computational biology, 12(6):e1004746, 2016.

[26] Y. Vander Meersche, G. Cretin, A. Gheeraert, J. C. Gelly, and T. Galochkina. Atlas: protein flexibility description from atomistic molecular dynamics simulations. Nucleic acids research, 52(D1):D384–D392, 2024.

[27] Alexander Rives, Joshua Meier, Tom Sercu, Siddharth Goyal, Zeming Lin, Jason Liu, Demi Guo, Myle Ott, C. Lawrence Zitnick, Jerry Ma, and Rob Fergus. Biological structure and function emerge from scaling unsupervised learning to 250 million protein sequences. Proceedings of the National Academy of Sciences, 118(15):e2016239118, 2021.

[28] R. Krishna, J. Wang, W. Ahern, P. Sturmfels, P. Venkatesh, I. Kalvet, et al. Generalized biomolecular modeling and design with rosettafold all-atom. Science, 384(6693):eadl2528, 2024.

[29] G. W. Kyro, T. Qiu, and V. S. Batista. A model-centric review of deep learning for protein design, 2025.

[30] D. Benson. Music: A Mathematical Offering. Cambridge University Press, 2006.

[31] G. Loy. Musimathics, Volume 1: The Mathematical Foundations of Music, volume 1. MIT Press, 2011.

[32] Ross D King and Colin G Angus. Pm—protein music. Bioinformatics, 12(3): 251–252, 1996.

[33] Rie Takahashi and Jeffrey H Miller. Conversion of amino-acid sequence in proteins to classical music: search for auditory patterns. Genome biology, 8(5): 405, 2007.

[34] Juliane Mössinger. The music of life, 2005.

[35] Setareh Ghavami, Hassan Toozandehjani, Ghazaleh Ghavami, and Soroush Sardari. Innovative protein translation into music and color image applicable for assessing protein alignment based on bio-mimicking human perception system. International Journal of Biological Macromolecules, 119: 896–901, 2018.

[36] J. Su and P. Zhou. Musical protein: Mapping the time sequence of music onto the spatial architecture of proteins. Computer Methods and Programs in Biomedicine, 252:108233, 2024.

[37] Irena Cosic. Macromolecular bioactivity: is it resonant interaction between macromolecules?-theory and applications. IEEE Transactions on Biomedical Engineering, 41(12): 1101–1114, 2002.

[38] Irena Cosic. The resonant recognition model of macromolecular bioactivity: theory and applications. Birkhäuser, 2012.

[39] David Eisenberg, Robert M Weiss, and Thomas C Terwilliger. The hydrophobic moment detects periodicity in protein hydrophobicity. Proceedings of the National Academy of Sciences, 81(1): 140–144, 1984.

[40] Irena Cosic. The resonant recognition model of bio-molecular interactions: possibility of electromagnetic resonance. Polish Journal of Medical Physics and Engineering, 7(1): 73–87, 2001.

[41] Irena Cosic and Elena Pirogova. Bioactive peptide design using the resonant recognition model. Nonlinear Biomedical Physics, 1(1): 7, 2007.

[42] Irena Cosic and Drasko Cosic. Dna-protein interactions at distance explained by the resonant recognition model. International Journal of Sciences, 13(11): 1–5, 2024.

[43] Felipe Bueno de Souza, Matheus Henrique Pimenta-Zanon, Dora Henriques, M Alice Pinto, Carlos Balsa, José Rufino, and Fabrício Martins Lopes. Resonant recognition model as a preprocessing technique for rna classification. In International Conference on Advanced Research in Technologies, Information, Innovation and Sustainability, pages 3–17. Springer, 2024.

[44] Charalambos Chrysostomou, Huseyin Seker, Nizamettin Aydin, and Parvez I Haris. Complex resonant recognition model in analysing influenza a virus subtype protein sequences. In Proceedings of the 10th IEEE International Conference on Information Technology and Applications in Biomedicine, pages 1–4. IEEE, 2010.

[45] N. A. O’Leary, M. W. Wright, J. R. Brister, S. Ciufo, D. Haddad, R. McVeigh, B. Rajput, B. Robbertse, B. Smith-White, D. Ako-Adjei, A. Astashyn, A. Badretdin, Y. Bao, O. Blinkova, V. Brover, V. Chetvernin, J. Choi, E. Cox, O. Ermolaeva, C. M. Farrell, T. Goldfarb, T. Gupta, D. Haft, E. Hatcher, W. Hlavina, V. S. Joardar, V. K. Kodali, W. Li, D. Maglott, P. Masterson, K. M. McGarvey, M. R. Murphy, K. O’Neill, S. Pujar, S. H. Rangwala, D. Rausch, L. D. Riddick, C. Schoch, A. Shkeda, S. S. Storz, H. Sun, F. Thibaud-Nissen, I. Tolstoy, R. E. Tully, A. R. Vatsan, C. Wallin, D. Webb, W. Wu, M. J. Landrum, A. Kimchi, T. Tatusova, M. DiCuccio, P. Kitts, T. D. Murphy, and K. D. Pruitt. Reference sequence (refseq) database at ncbi: current status, taxonomic expansion, and functional annotation. Nucleic Acids Research, 44:D733–D745, 2016.

[46] The UniProt Consortium. Uniprot: the universal protein knowledgebase in 2023. Nucleic Acids Research, 51:D523–D531, 2023.

[47] G. Wang, C. Schmidt, X. Li, and Z. Wang. APD6: the antimicrobial peptide database is expanded to promote research and development by deploying an unprecedented information pipeline. Nucleic Acids Research, page gkaf860, 2025.

[48] M. H. Saier Jr, V. S. Reddy, G. Moreno-Hagelsieb, K. J. Hendargo, Y. Zhang, V. Iddamsetty, and A. Medrano-Soto. The transporter classification database (TCDB): 2021 update. Nucleic acids research, 49(D1):D461–D467, 2021.

[49] J. Yang, A. Mora, S. Liu, B. Wittmann, A. Anandkumar, F. Arnold, and Y. Yue. Care: a benchmark suite for the classification and retrieval of enzymes. In Advances in Neural Information Processing Systems, volume 37, pages 3094–3121, 2024.

[50] W. Li and A. Godzik. Cd-hit: a fast program for clustering and comparing large sets of protein or nucleotide sequences. Bioinformatics, 22(13): 1658–1659, 2006.

[51] N. Japkowicz and S. Stephen. The class imbalance problem: A systematic study. In Intelligent Data Analysis, volume 6, pages 429–449, 2002.

[52] Giovanni Bellesia, Andrew Iain Jewett, and Joan-Emma Shea. Sequence periodicity and secondary structure propensity in model proteins. Protein Science, 19(1): 141–154, 2010.

[53] C Spence. Crossmodal correspondences: A tutorial review. Attention, Perception, & Psychophysics, 73(4):971–995, 2011.

[54] A. Kuzmanic and B. Zagrovic. Determination of ensemble-average pairwise root mean-square deviation from experimental b-factors. Biophysical journal, 98(5): 861–871, 2010.

[55] Zhuang Liu, Hanzi Mao, Chao-Yuan Wu, Christoph Feichtenhofer, Trevor Darrell, and Saining Xie. A convnet for the 2020s. In Proceedings of the IEEE/CVF Conference on Computer Vision and Pattern Recognition, pages 11976–11986, 2022.

[56] Facebook AI. Model repository for esm2 t6 8m ur50d. https://huggingface.co/facebook/esm2_t6_8M_UR50D, 2022. Accessed: 2025-08-26.

[57] Facebook AI. Model repository for esm2 t12 35m ur50d. https://huggingface.co/facebook/esm2_t12_35M_UR50D, 2022. Accessed: 2025-08-26.

[58] Mark Johnson, Irena Zaretskaya, Yan Raytselis, Yuri Merezhuk, Scott McGinnis, and Thomas L Madden. NCBI BLAST: a better web interface. Nucleic acids research, 36(Web Server issue):W5–W9, 2008. Accessed: 2025-08-26.

[59] Thomas Cover and Peter Hart. Nearest neighbor pattern classification. IEEE transactions on information theory, 13(1): 21–27, 1967.

[60] K. S. Sarkisyan, D. A. Bolotin, M. V. Meer, D. R. Usmanova, A. S. Mishin, G. V. Sharonov, and F. A. Kondrashov. Local fitness landscape of the green fluorescent protein. Nature, 533(7603): 397–401, 2016.

[61] BFD Datasets. Bfd. https://bfd.mmseqs.com/, 2018. Accessed: 2025-10-01.

[62] nferruz. Ur50 2021 04. https://huggingface.co/datasets/nferruz/UR50_2021_04, 2022. Accessed: 2025-08-26.

[63] E. Brattico and M. Delussi. Making sense of music: Insights from neurophysiology and connectivity analyses in naturalistic listening conditions. Hearing Research, 441:108923, 2024.

[64] D. Bountouridis, D. G. Brown, F. Wiering, and R. C. Veltkamp. Melodic similarity and applications using biologically-inspired techniques. Applied Sciences, 7(12): 1242, 2017.

[65] T. Giesa, D. I. Spivak, and M. J. Buehler. Reoccurring patterns in hierarchical protein materials and music: the power of analogies. BioNanoScience, 1: 153–161, 2011.

[66] J. Y. Wong, J. McDonald, M. Taylor-Pinney, D. I. Spivak, D. L. Kaplan, and M. J. Buehler. Materials by design: Merging proteins and music. Nano Today, 7(6): 488–495, 2012.

[67] H. Dong, J. Xu, Y. Yang, R. Zhao, S. Wu, C. Yuan, …, and L. Han. MeGraph: capturing long-range interactions by alternating local and hierarchical aggregation on multi-scaled graph hierarchy. In Advances in Neural Information Processing Systems, volume 36, pages 63609–63641, 2023.

[68] Y. Wang, C. Cheng, S. Li, Y. Ren, B. Shao, G. Liu, …, and N. Zheng. Neural P3M: A Long-Range Interaction Modeling Enhancer for Geometric GNNs. In Advances in Neural Information Processing Systems, volume 37, pages 120336–120365, 2024.

[69] Ashish Vaswani, Noam Shazeer, Niki Parmar, Jakob Uszkoreit, Llion Jones, Aidan N Gomez, Łukasz Kaiser, and Illia Polosukhin. Attention is all you need. Advances in neural information processing systems, 30, 2017.

[70] G. Amitai, A. Shemesh, E. Sitbon, M. Shklar, D. Netanely, I. Venger, and S. Pietrokovski. Network analysis of protein structures identifies functional residues. Journal of Molecular Biology, 344(4): 1135–1146, 2004.

[71] R. Braun, M. Tfirn, and R. M. Ford. Listening to life: Sonification for enhancing discovery in biological research. Biotechnology and Bioengineering, 121(10): 3009–3019, 2024.

[72] E. J. Martin, T. R. Meagher, and D. Barker. Using sound to understand protein sequence data: new sonification algorithms for protein sequences and multiple sequence alignments. BMC Bioinformatics, 22: 1–17, 2021.

[73] K. Patil, D. Pressnitzer, S. Shamma, and M. Elhilali. Music in our ears: the biological bases of musical timbre perception. PLoS computational biology, 8(11):e1002759, 2012.

[74] Yuan Gong, Yu-An Chung, and James Glass. Ast: Audio spectrogram transformer. arXiv preprint arXiv:2104.01778, 2021.

[75] S. Orouji, M. C. Liu, T. Korem, and M. A. Peters. Domain adaptation in small-scale and heterogeneous biological datasets. Science Advances, 10(51):eadp6040, 2024.

[76] H. W. Hellinga. Rational protein design: combining theory and experiment. Proceedings of the National Academy of Sciences, 94(19): 10015–10017, 1997.

[77] T. Schmidt, A. Bergner, and T. Schwede. Modelling three-dimensional protein structures for applications in drug design. Drug discovery today, 19(7): 890–897, 2014.

[78] N. W. Tay, F. Liu, C. Wang, H. Zhang, P. Zhang, and Y. Z. Chen. Protein music of enhanced musicality by music style guided exploration of diverse amino acid properties. Heliyon, 7(9), 2021.

