## Appendix for "From Signal to Symphony: Exploring 2D Sequence Representations for Protein Function Prediction"

### A Data and Sources

The primary data for this study were sourced from several publicly available databases. Table S1 lists the types of materials and their respective sources.

Table S1: Materials and Sources for the Study.

| Types of Materials | Sources | URLs |
|---|---|---|
| Protein FASTA Sequences | NCBI Database | https://www.ncbi.nlm.nih.gov/ |
| | UniProt Database | https://www.unipro.org/ |
| | Antimicrobial Peptide Database | https://aps.unmc.edu/ |
| | Transporter Classification Database | https://www.tcdb.org/ |
| | CARE database | https://github.com/jsunn-y/CARE/ |

### B Quantitative Protein Sonification Framework

#### B.1 Objective and Principle

This section details the quantitative model used to translate protein sequences into musical feature representations. The goal is to create a reproducible, rule-based system for generating 2D spectrograms from 1D sequence data, providing a foundation for the machine learning analyses presented in the main text.

Our mapping framework is built on the principle that key physicochemical properties of amino acids can be systematically encoded into distinct musical parameters. **The quantitative model implemented and tested in this study focuses exclusively on these local amino acid properties.** In Section B.3, we also discuss a conceptual framework for how this model could be extended in future work to incorporate higher-order structural information.

#### B.2 Quantitative Mapping Rules Implemented in This Study

##### B.2.1 Mapping of Amino Acid Physicochemical Properties

The primary translation step converts the linear sequence of amino acids into a sequence of musical notes, where each note's properties are determined by the corresponding amino acid's physicochemical characteristics.

**Hydrophobicity to Pitch Mapping.** We used the Kyte-Doolittle hydrophobicity scale to define the pitch of each note. The mapping function $f_{\text{pitch}}$ translates an amino acid's hydrophobicity index $H$ into a MIDI note number.

$$Pitch_{MIDI} = \text{round}(Pitch_{base} \pm (H \times S_{\text{pitch}})) \tag{1}$$

where $Pitch_{MIDI}$ is the final MIDI note value, $Pitch_{base}$ is a reference pitch (set to 60 for Middle C), $H$ is the hydrophobicity index, and $S_{\text{pitch}}$ is a scaling factor (set to 3). The sign ($\pm$) is determined by the specific mapping scheme being tested (e.g., '-' for the theory-driven model, '+' for the inverted-semantics model, as described in the main text).

**Polarity to Timbre Mapping.** Amino acid polarity determines the timbre (tonal quality) of the note. We used a binary mapping based on a polarity threshold $P_{\text{threshold}}$.

$$\text{Timbre} = f_{\text{timbre}}(\text{aa}) = \begin{cases} \text{Preset A (e.g., Piano)} & \text{if } P(\text{aa}) > P_{\text{threshold}} \text{ (Polar)} \\ \text{Preset B (e.g., Cello)} & \text{otherwise (Non-polar)} \end{cases} \tag{2}$$

In our implementation, Preset A is characterized by a "bright" harmonic spectrum (more high-frequency overtones), while Preset B is "darker" (emphasizing lower-frequency fundamentals and harmonics).

#### B.3 Conceptual Framework for Future Extensions: Mapping Higher-Order Structures

While the quantitative model evaluated in this study is intentionally streamlined to focus on first principles (i.e., amino acid properties), the sonification framework offers a rich conceptual space for future extensions by incorporating higher-order structural information. The following outlines potential mapping strategies that could be explored in subsequent work to create even more nuanced representations. **Note: These conceptual rules were not implemented or tested in the current study but are presented here to illustrate the framework's potential.**

###### Potential Mapping of Secondary Structure

One could map predicted or known secondary structures to distinct musical motifs to make them audibly recognizable:

- **$\alpha$-helix:** Could be represented by smooth, connected (legato) melodic contours, reflecting its regular, continuous helical structure.
- **$\beta$-sheet:** Could be mapped to more detached (staccato) or repetitive chordal patterns, evoking its rigid, pleated nature.

###### Potential Mapping of Tertiary and Quaternary Structure

Global structural features could be mapped to larger-scale musical forms:

- **Global Fold:** The overall protein architecture could inform the musical form. For instance, a globular protein might inspire a compact, recurring musical theme (a rondo), while a fibrous protein could be translated into a more linear, through-composed piece.
- **Multimeric Complexes:** Polyphonic techniques offer a natural analogy for protein complexes. Each subunit could be conceptualized as an independent melodic line (a voice in counterpoint), with their musical interplay designed to reflect the geometric and functional relationships between the subunits.

#### B.4 Technical Implementation Details

This section provides the specific implementation details for the Quantitative Sonification Framework. All parameters and mapping rules are explicitly defined here to ensure full transparency and reproducibility of our methods.

##### B.4.1 Amino Acid to Musical Parameter Mapping Rules

The core of the framework is a set of deterministic rules that convert biophysical properties of each amino acid into musical parameters.

The pitch of a note is derived from the Kyte-Doolittle hydrophobicity scale. The formula used is:

$$\texttt{MIDI Pitch} = \text{round}(\texttt{BASE\_NOTE} - (\texttt{Kyte-Doolittle\_Value} \times \texttt{SCALING\_FACTOR})) \tag{3}$$

Where the base pitch `BASE_NOTE` is set to 60 (MIDI for Middle C) and the `SCALING_FACTOR` is 3.0. The final calculated pitch is clipped to the valid MIDI range of [0, 127]. The complete mapping is detailed in Table S2.

**Timbre and Polarity Mapping.** The instrument timbre is determined by the amino acid's polarity, distinguishing residues likely to be on the protein's surface from those in its core.

- **Polarity Scale:** We use the Zimmerman polarity scale. The specific values are: A: 8.1, R: 10.5, N: 11.6, D: 13.0, C: 5.5, Q: 10.5, E: 12.3, G: 9.0, H: 10.4, I: 5.2, L: 4.9, K: 11.3, M: 5.3, F: 5.2, P: 8.0, S: 9.2, T: 8.6, W: 5.4, Y: 6.2, V: 5.9.
- **Mapping Rule:** A threshold is set at the median of these values (`POLARITY_THRESHOLD` = 8.8).
- **Timbre Implementation:**
  - **Non-polar** (Polarity $\leq$ 8.8) amino acids are mapped to a "bright" timbre: **Acoustic Grand Piano** (General MIDI Program Change number **0**).
  - **Polar** (Polarity > 8.8) amino acids are mapped to a "dark" timbre: **Cello** (General MIDI Program Change number **42**).

Table S2: Complete mapping of amino acids to Kyte-Doolittle hydrophobicity values and the resulting calculated MIDI pitch.

| Amino Acid | Kyte-Doolittle Value | Calculated MIDI Pitch |
|---|---|---|
| Alanine (A) | 1.8 | 55 |
| Arginine (R) | -4.5 | 74 |
| Asparagine (N) | -3.5 | 71 |
| Aspartic Acid (D) | -3.5 | 71 |
| Cysteine (C) | 2.5 | 52 |
| Glutamine (Q) | -3.5 | 71 |
| Glutamic Acid (E) | -3.5 | 71 |
| Glycine (G) | -0.4 | 61 |
| Histidine (H) | -3.2 | 70 |
| Isoleucine (I) | 4.5 | 46 |
| Leucine (L) | 3.8 | 49 |
| Lysine (K) | -3.9 | 72 |
| Methionine (M) | 1.9 | 54 |
| Phenylalanine (F) | 2.8 | 52 |
| Proline (P) | -1.6 | 65 |
| Serine (S) | -0.8 | 62 |
| Threonine (T) | -0.7 | 62 |
| Tryptophan (W) | -0.9 | 63 |
| Tyrosine (Y) | -1.3 | 64 |
| Valine (V) | 4.2 | 47 |

**Other Musical Parameters.**

- **Note Duration:** The duration of each note is dynamically determined by the molecular weight of the corresponding amino acid, creating a natural rhythm based on residue size. The formula is:

$$\texttt{Duration (in beats)} = \frac{\texttt{Molecular\_Weight}}{100.0} \tag{4}$$

  The tempo is set to a constant 120 beats per minute.

- **Note Velocity (Volume):** The velocity for all notes is fixed at a constant value of **100** on the MIDI scale of 0–127. This ensures that volume does not act as a confounding variable in the spectral representation.

##### B.4.2 Software Libraries and Tools

**MIDI File Generation.** We used the Python library `midiutil` to programmatically create standard MIDI files (`.mid`) from the musical parameters generated by our mapping rules.

**Audio Rendering (MIDI-to-WAV).**

- **Synthesizer:** The open-source software synthesizer **FluidSynth** was used to render the MIDI files into audio waveforms. This process was automated using the Python wrapper library `midi2audio`.

- **SoundFont Library:** The **GeneralUser GS v1.471** SoundFont (`GeneralUser.sf2`) was used as the source for all instrument sounds. It is available at https://musical-artifacts.com/artifacts/4625 or from our GitHub repository at https://github.com/wyqmath/Symphony_of_Fate. This is a comprehensive General MIDI (GM) compatible sound bank. The output audio was saved as WAV files with a sample rate of 16,000 Hz.

**Spectrogram Generation Parameters.** The final 2D representations (Mel spectrograms) were generated from the WAV audio files using the Python library `librosa`. The specific parameters for the `librosa.feature.melspectrogram` function were as follows, ensuring a standardized transformation:

- **FFT Window Size (`n_fft`):** 2048 samples. This determines the frequency resolution of the analysis.
- **Hop Length (`hop_length`):** 512 samples. This sets the time-domain resolution by defining the step size between consecutive analysis windows.
- **Number of Mel Bands (`n_mels`):** 128. This defines the vertical resolution (number of frequency bins) of the final spectrogram.
- **Window Type (`window`):** 'hann'. The Hann window function was applied to each frame to reduce spectral leakage. This is the default setting in `librosa`.
- **Frequency Range (`fmin`, `fmax`):** Default values were used, spanning from 0 Hz to the Nyquist frequency (`sr`/2.0).

##### B.5 Justification for the Chosen Mapping Scheme and Discussion of Alternatives

We acknowledge that any specific mapping involves a choice. The experiments in the main text (see Results, Table 2) were designed to rigorously test the impact of our core, implemented mapping rules. **The conceptual ideas for mapping higher-order structures, outlined in Section B.3, represent avenues for future research and were deliberately excluded from the current quantitative model to allow for a clearer, more controlled investigation into the foundational principles of the 1D-to-2D transformation.**

#### C Biophysical Validation: Molecular Dynamics Simulation

To provide a dynamic context for protein flexibility, which was subsequently compared with experimental B-factors, we performed all-atom molecular dynamics (MD) simulations for each protein. All simulations were conducted using the GROMACS 2025.2. The following protocol details the standardized and reproducible workflow employed.

##### C.1 System Preparation

**PDB Structure Pre-processing.** The initial crystal structure for each protein was downloaded from the Protein Data Bank (PDB). To ensure a stable and complete starting structure for the simulation, the raw PDB file was pre-processed using PDBFixer. This step involved: (1) replacing non-standard amino acid residues with their standard equivalents, (2) adding missing heavy atoms within residues, (3) modeling missing residues and loops where applicable, and (4) removing all water molecules and non-essential heteroatoms (e.g., ligands, crystallization agents).

**Protonation and Force Field Assignment.** Hydrogen atoms were added to the cleaned structure, and the protonation states of ionizable residues (such as Aspartic Acid, Glutamic Acid, Lysine, Arginine, and Histidine) were determined assuming a physiological pH of 7.0, a functionality also handled by PDBFixer. The system was then prepared for simulation using the GROMACS `pdb2gmx` tool with the **AMBER99SB-ILDN** force field and the **TIP3P** water model.

**Solvation and Ionization.** The protein was centered in a cubic simulation box, ensuring a minimum distance of 1.0 nm between any protein atom and the box boundaries. The box was subsequently filled with TIP3P water molecules. To neutralize the system's net charge and mimic physiological ionic strength, sodium ($Na^+$) and chloride ($Cl^-$) ions were added to achieve a final salt concentration of 0.15 M.

##### C.2 MD Simulation Parameters

The simulation proceeded through three main stages: energy minimization, system equilibration, and production MD. All parameter files (`.mdp`) are detailed below.

Table S3: Conceptual Space of Protein-to-Music Mapping Rules.

| **Protein Feature** | **Musical Element** | **Mapping Rule Used in This Study** | **Considered Alternative(s)** | **Rationale for Alternatives** |
|---|---|---|---|---|
| **Hydrophobicity** | Pitch | **High Hydrophobicity → Low Pitch** (Rationale: Analogy to energetic stability of the buried core and psychoacoustically "grounded" low frequencies.) | **High Hydrophobicity → High Pitch** (This was implemented as our 'Inverted Semantics' control.) | Analogy to high tension or energetic strain within the tightly packed protein core. |
| **Structural Flexibility (B-factor)** | Rhythm | **High Flexibility → Complex/Syncopated Rhythm** (Rationale: Dynamic, irregular motion maps to rhythmically complex patterns.) | **1.** High Flexibility → Faster Tempo<br>**2.** High Flexibility → Increased Vibrato/Tremolo | 1. A direct mapping of faster atomic motion to a faster musical tempo.<br>2. Representing structural fluctuation as modulation of pitch (vibrato) or amplitude (tremolo). |
| **Secondary Structure** | Harmony / Timbre | **$\alpha$-helix → Smooth Melodic Contour**<br>**$\beta$-sheet → Repetitive Chords** | **$\alpha$-helix / $\beta$-sheet → Distinct Instrument Timbres** (e.g., $\alpha$-helix = Flute; $\beta$-sheet = Oboe) | Mapping unique structural motifs to unique sonic colors (timbres) to make them perceptually distinct. |
| **Polarity** | Timbre | **Polar → "Bright" Timbre**<br>**Non-polar → "Dark" Timbre** | **Polarity → Harmonic Consonance/Dissonance** | Mapping stable surface interactions (polar residues) to harmonically stable sounds (consonance). |

**Energy Minimization.** To remove any steric clashes or unfavorable geometries introduced during system preparation, an energy minimization was performed.

- **Algorithm:** Steepest Descent.
- **Maximum Steps:** 50,000.
- **Convergence Threshold:** The minimization proceeded until the maximum force on any atom was less than 1000.0 kJ $mol^{-1}$ $nm^{-1}$.

**System Equilibration.** A two-phase equilibration was conducted to gradually bring the system to the desired temperature and pressure. During both phases, position restraints with a force constant of 1000 kJ $mol^{-1}$ $nm^{-2}$ were applied to all protein heavy atoms to allow the solvent to relax around the protein.

- **NVT Equilibration (Constant Volume):** The system was heated to 300 K over 100 ps. Temperature was controlled using the V-rescale thermostat with a coupling time constant of 0.1 ps. Separate coupling groups were used for the protein and non-protein (solvent and ions) components.
- **NPT Equilibration (Constant Pressure):** The system was equilibrated for an additional 100 ps at a constant pressure of 1.0 bar. Pressure was maintained using the Parrinello-Rahman barostat with a coupling time constant of 2.0 ps. Temperature coupling was identical to the NVT phase.

**Production MD.** Following equilibration, the position restraints were removed, and a 100 ns production simulation was carried out for data collection.

- **Integrator:** Leap-frog MD integrator with a time step of 2 fs.
- **Constraints:** All bonds involving hydrogen atoms were constrained using the LINCS algorithm.
- **Non-bonded Interactions:** A cut-off distance of 1.0 nm was used for short-range van der Waals and Coulomb interactions. Long-range electrostatic interactions were handled using the Particle Mesh Ewald (PME) method.
- **Thermostat and Barostat:** Temperature (300 K) and pressure (1.0 bar) were maintained using the same V-rescale and Parrinello-Rahman schemes as in the NPT equilibration phase.

##### C.3 Trajectory Analysis

**Trajectory Pre-processing.** To calculate meaningful Root Mean Square Fluctuation (RMSF) values, the raw production trajectory was first corrected. The `gmx trjconv` tool was used to correct for periodic boundary conditions by ensuring whole molecules were not split across box boundaries. Subsequently, the trajectory was aligned by performing a least-squares fit of the protein backbone atoms to the backbone of the equilibrated starting structure. This critical step removes global translational and rotational movements, ensuring that the calculated RMSF reflects true internal atomic fluctuations.

**RMSF Calculation.** The RMSF for each residue was calculated based on the C$\alpha$ atom positions over the entire 100 ns production trajectory. This was accomplished using the GROMACS analysis tool `gmx rmsf`. The specific command used was:

```
gmx rmsf -s md_production.tpr -f md_fit.xtc -o rmsf.xvg -res
```

where `md_fit.xtc` is the pre-processed trajectory. The output `rmsf.xvg` file, containing RMSF values in nm per residue, was then used for direct comparison with the experimental B-factors.

#### D Implementation of Classification and Prediction Systems

This study implements and compares multiple classification pipelines to demonstrate the progressive benefit of evolving data representation and model complexity. The implementation details for our primary end-to-end fusion model and all benchmark models are provided below.

#### D.1 End-to-End Fusion Model with Two-Phase Training

Our primary model is a sophisticated, end-to-end deep learning system trained in two distinct phases to maximize performance by effectively leveraging pre-trained knowledge and fusing visual and acoustic features.

- **Data Augmentation:** A comprehensive augmentation pipeline was applied to the training spectrograms using the `timm` library. Transformations included resizing to the model's expected input size, random rotations (10 degrees), random resized crops, horizontal flips, and random erasing. Crucially, we also applied **SpecAugment**, a technique that randomly masks blocks of frequency (up to 25 bins) and time (up to 50 steps) information, forcing the model to learn more robust and generalized features.

- **Phase 1: Image-Only Backbone Pre-training.**
  - **Objective:** To fine-tune a powerful image recognition model on our spectrogram data distribution. This generates a set of highly relevant initial weights for the visual branch of the final fusion model, adapting the ImageNet-trained model to our specific domain.
  - **Architecture:** An `ImageOnlyModel` was constructed, consisting of a `ConvNeXt-Tiny` backbone (pre-trained on ImageNet, with input channels modified to 1 for grayscale) and a classification head containing a linear layer, ReLU activation, and dropout.
  - **Training:** The model was trained on spectrogram images alone, using the AdamW optimizer with a cosine annealing learning rate schedule and label smoothing (0.1). Early stopping (patience=15) was used to save the weights corresponding to the best validation loss.

- **Phase 2: Attention Fusion Model Training.**
  - **Architecture:** The final `AttentionFusionModel` was constructed with a dual-branch architecture:
    - **Visual Branch:** A `ConvNeXt-Tiny` backbone, with its weights initialized from the model saved in Phase 1.
    - **Acoustic Branch:** This branch processes the temporal dimension of the audio. For each spectrogram, a sequence of 60 Mel-Frequency Cepstral Coefficients (MFCCs) per time step is extracted. This sequence is fed into a Gated Recurrent Unit (GRU) block, followed by an attention mechanism. The specific parameters are:
      - **GRU:** The GRU has a hidden size of **128**, consists of **2 layers**, and is **bidirectional**. The output dimension for the sequence is therefore 256 (128 hidden size $\times$ 2 directions).
      - **Attention Mechanism:** A feed-forward based additive attention mechanism is applied to the GRU's output sequence. This mechanism uses a two-layer neural network to compute alignment scores. The first linear layer reduces the feature dimension from 256 to 128, followed by a Tanh activation. The second linear layer projects the 128-dimensional vector to a single scalar score for each time step. These scores are then normalized via Softmax to produce the final attention weights.
    - **Fusion and Classification:** The feature vector from the visual backbone and the context vector from the attentive acoustic branch are concatenated and passed to a final classification head for prediction.
  - **Training Strategy:** A staged fine-tuning approach was used. For the first 30 epochs, the visual backbone was frozen, and only the newly added components (the acoustic branch and the final classifier) were trained. Subsequently, the entire network was unfrozen and fine-tuned end-to-end. We used the AdamW optimizer with a Cosine Annealing learning rate schedule and **differential learning rates**: the pre-trained backbone was updated with a smaller learning rate ($1 \times 10^{-5}$) than the newly initialized head components ($1 \times 10^{-4}$). Early stopping (patience=15) was used to prevent overfitting.

#### D.2 Benchmark Model Implementations

To contextualize the performance of our proposed fusion model, we implemented and evaluated several distinct benchmark systems.

##### D.2.1 Classical ML with Spectral Features

This model establishes a classical machine learning baseline using spectral features derived directly from the 1D physicochemical representation of protein sequences.

- **Signal Conversion and Feature Extraction:** Each protein sequence was first converted into a 1D numerical signal using the Kyte-Doolittle hydrophobicity scale. A Fast Fourier Transform (FFT) was then applied to this signal, and the resulting power spectrum was used as the feature vector for each sequence.
- **Data Augmentation:** To increase the robustness of the model, the training set was augmented using three techniques: (1) conservative mutation based on the BLOSUM62 matrix, (2) random cropping of sequences to 90% of their original length, and (3) the addition of a small amount of Gaussian noise to the hydrophobicity signal before the FFT.
- **Feature Selection:** A two-step feature selection process was employed to reduce dimensionality and select the most informative spectral components. First, `SelectFromModel` with a `RandomForestClassifier` was used for broad feature filtering. Second, Recursive Feature Elimination (RFE) was applied to select the top 50 features.
- **Model Architecture:** The final classifier was a soft-voting ensemble (`VotingClassifier`) combining four models: a `RandomForestClassifier`, an `XGBClassifier`, a Support Vector Classifier (`SVC`), and a Multi-layer Perceptron (`MLPClassifier`).
- **Training:** The model was evaluated using 5-fold stratified cross-validation and trained on the full augmented training set to produce the final predictions.

##### D.2.2 1D CNN Baseline

This model was designed to establish a strong deep learning baseline using a one-dimensional data representation.

- **Signal Conversion:** Each protein sequence was converted into a 1D numerical signal using the Kyte-Doolittle hydrophobicity scale.
- **Data Augmentation:** The training set was augmented via conservative mutation, random cropping (90% subsequence), and the addition of a small amount of random noise (mean=0, std=0.05) to the hydrophobicity signal. All signals were padded to a uniform maximum length determined from the training set.
- **Model Architecture:** A 1D CNN was implemented in PyTorch, consisting of three sequential convolutional blocks followed by a fully connected head. The detailed parameters for each convolutional layer are provided in Table S4.
- **Training:** The model was trained using the Adam optimizer (LR=0.001, weight decay=1e-4) with a Cross-Entropy Loss function and an early stopping strategy (patience=15) based on validation loss.

Table S4: Detailed architecture of the 1D Convolutional Neural Network. Each convolutional layer is followed by Batch Normalization, a ReLU activation, Max Pooling (kernel size 2, stride 2), and Dropout.

| Layer | In Channels | Out Channels | Kernel Size | Stride | Padding |
|---|---|---|---|---|---|
| Convolutional Block 1 | 1 | 64 | 7 | 1 | 3 |
| Convolutional Block 2 | 64 | 128 | 5 | 1 | 2 |
| Convolutional Block 3 | 128 | 256 | 3 | 1 | 1 |

##### D.2.3 Protein Language Model Benchmarks

We benchmarked against state-of-the-art transformer-based Protein Language Models (PLMs).

**ESM-2 Benchmark.** We evaluated two model sizes from the ESM-2 family, `esm2_t6_8M_UR50D` (8M parameters) and `esm2_t12_35M_UR50D` (35M parameters).

- **Scenarios:** Two distinct training scenarios were tested:
    1. **Fine-tuning:** The official pre-trained ESM-2 models were used, and only a linear classification head was fine-tuned on our training data.
    2. **From Scratch:** The entire ESM-2 architecture was trained solely on our dataset without pre-trained weights to directly assess data efficiency.
- **Data Preparation:** Sequences were tokenized using the official ESM-2 tokenizer and padded or truncated to a maximum length of 1024 tokens.
- **Model Architecture:** The classifier consisted of the base ESM-2 model followed by a classification head that takes the final hidden state of the special `[CLS]` token for prediction.
- **Training:** The model was trained using the AdamW optimizer (LR=1e-5, weight decay=0.01) with early stopping (patience=15).

**ProtBERT Benchmark.** In addition to ESM-2, we benchmarked against the **ProtBERT** model (`Rostlab/prot_bert`).

- **Scenario:** We evaluated the model in a **fine-tuning** setting, leveraging its pre-trained weights learned from a massive corpus of protein sequences.
- **Data Preparation:** A specific preprocessing step was required for the ProtBERT tokenizer, where spaces were inserted between each amino acid in the sequence. The sequences were then tokenized and padded or truncated to a maximum length of 1024 tokens.
- **Model Architecture:** The classifier architecture consisted of the pre-trained ProtBERT base model, followed by a dropout layer (p=0.3) and a single linear layer for the final classification. The input to the classifier is the representation of the `[CLS]` token.
- **Training:** The model was fine-tuned using the AdamW optimizer (LR=1e-5, weight decay=0.01) with a Cross-Entropy Loss function. Training was regularized with an early stopping mechanism (patience=15) based on validation loss.

###### D.2.4 Homology-Based Benchmarks

To provide a robust comparison against established bioinformatics methods, two distinct homology-based benchmarks were implemented. Both rely on the BLAST algorithm but use different databases and classification strategies. In both cases, the ranking of homologous sequences (neighbors) is determined by BLAST's default sorting algorithm, which prioritizes hits with the lowest **E-value**, using the bit-score as a secondary criterion for tie-breaking.

**Benchmark 1: kNN Classification using an Internal BLAST Database.** This benchmark serves as a classical k-Nearest Neighbors (kNN) baseline, where classification relies solely on the information contained within our dataset.

- **Database:** A custom BLAST database was constructed exclusively from the protein sequences in our designated **training set**.
- **BLAST Search:** For each sequence in the test set, a `blastp` search was performed against this internal database. The command-line template was:

```
blastp -query <query_file> -db <training_set_db> -evalue 1e-5 \
-max_target_seqs 5 -outfmt '6 sseqid bitscore'
```

- **Classification Logic:** The functional class was assigned based on a simple majority vote among the top 5 nearest neighbors ($k = 5$) identified by the BLAST search. The class labels of the neighbors were retrieved from the training set annotations. If no significant hits were found, a class was assigned randomly.

**Benchmark 2: Hierarchical Classification using the Swiss-Prot Database.** This benchmark was designed to emulate the "gold standard" approach of identifying homologs in a comprehensive, manually curated public database. It employs a more sophisticated, rule-based classification logic to maximize precision.

- **Database:** A local copy of the UniProtKB/Swiss-Prot database (**Release 2025_03**) was used.
- **BLAST Search:** The search was parallelized externally using Python's `multiprocessing` library. The command-line template for each search was:

```
blastp -query <query_file> -db <path_to_swissprot_db> -evalue 1e-5 \
-max_target_seqs 5 -outfmt '6 stitle pident qcovs'
```

- **Hierarchical Classification Logic:** A multi-step process was applied to simulate expert manual annotation and filtering:
  1. **Keyword-based Class Mapping:** A comprehensive mapping system was developed to assign a functional class based on the text description (title) of a BLAST hit. This system used prioritized keyword lists, including highly specific terms (e.g., "collagen", "myosin") for high-confidence assignment and more general terms (e.g., "enzyme", "transporter"), as well as negative keywords to resolve ambiguity.
  2. **High-Confidence Top Hit Analysis:** The top-ranked hit was analyzed first. Its title was mapped to a functional class using the keyword system. A prediction was made only if the percentage identity (`pident`) met a predefined threshold.
  3. **Identity Thresholding:** A general minimum identity of **30%** was required for a hit to be considered for classification. Sequences with a top hit below this threshold were immediately marked as 'Unclassified'. To enhance sensitivity for functionally diverse families, this threshold was adaptively lowered for specific classes (e.g., to 20% for Enzymes and 15% for Antimicrobial peptides).
  4. **Fallback to Majority Vote:** If the top hit did not yield a high-confidence classification (e.g., failed the identity threshold), the titles of all top 5 hits were mapped to functional classes. The final prediction was then determined by a majority vote among these 5 potential classes.
  5. **Final Assignment:** If neither step 2 nor step 4 yielded a definitive class, the query sequence was labeled as 'Unclassified'.

### E Implementation of Generative Model for GFP Variants

This section details the computational framework for generating and selecting novel Green Fluorescent Protein (GFP) variants, as presented in the main text. Our approach implements an *in silico* directed evolution loop, where variant generation is achieved through a guided mutation-and-selection process rather than a deep generative model like a diffusion model or GAN. The loop is driven by two key components: a high-performance fitness predictor and a novel "Harmonic Score" derived from our protein sonification framework. This dual-objective strategy aims to identify variants that are not only predicted to be highly fluorescent but also possess desirable structural or aesthetic properties as captured by the harmonic score.

#### E.1 Fitness Prediction Model

A supervised learning model was trained to predict the fluorescence fitness of a given GFP sequence. This model serves as the primary oracle for guiding the selection process.

##### E.1.1 Model Architecture

The fitness predictor is a regression model built on top of a pre-trained protein language model. The architecture consists of:

1. **Backbone:** The `esm2_t6_8M_UR50D` model from the ESM-2 family, a transformer-based model with 8 million parameters, pre-trained on a massive corpus of protein sequences.
2. **Regression Head:** The sequence embedding is obtained by taking the mean of the final hidden states of the ESM-2 model across all amino acid positions (masked for padding). This embedding is then passed through a two-layer Multi-Layer Perceptron (MLP) to predict a single scalar fitness value. The MLP architecture is:
   - A linear layer mapping the ESM-2 hidden size (320) to a hidden dimension of 128.
   - A Rectified Linear Unit (ReLU) activation function.
   - A final linear layer mapping the 128-dimensional vector to a single output neuron.

This architecture is implemented using PyTorch and the Hugging Face Transformers library.

###### E.1.2 Training Data

- **Data Source:** The training data was derived from the experimental fluorescence dataset of GFP variants published by Sarkisyan et al., which originally contains 54,025 sequences.
- **Data Filtering and Size:** To ensure the model was trained on functionally relevant data and to remove noisy, low-fluorescence entries, we applied a preprocessing filter. Only variants with a log fluorescence intensity greater than a threshold of 1.35 were retained for training. This filtering process resulted in a final high-quality training set of **41,745** sequences.
- **Preprocessing:** The target variable, log fluorescence intensity (referred to as `conv(%)` in our implementation), for the filtered dataset was normalized to a range of [0, 1] using a `MinMaxScaler`. Protein sequences were tokenized using the standard ESM-2 tokenizer.

###### E.1.3 Training Details

The fitness predictor was trained with the following hyperparameters:

- **Epochs:** 50
- **Batch Size:** 256 per GPU
- **Optimizer:** Adam, with a learning rate of $1 \times 10^{-4}$.
- **Loss Function:** Mean Squared Error (MSE), suitable for the regression task.

The model with the lowest validation loss was saved and used for all subsequent fitness predictions.

##### E.2 Harmonic Score Calculation

The "Harmonic Score" is a quantitative metric designed to capture the tonal stability and consonance of a protein sequence when translated into music via our sonification framework. This score serves as a secondary, orthogonal objective for variant selection. The calculation follows a three-step, fully deterministic process:

1. **Sequence-to-Music Sonification:** Each amino acid in a protein sequence is mapped to a musical note with specific properties based on its physicochemical characteristics:
   - **Pitch:** Determined by the Kyte-Doolittle hydrophobicity scale. The MIDI pitch value is calculated using the formula:
   $$Pitch_{MIDI} = \text{round}(60 - (H_{\text{KD}} \times 3.0)) \tag{5}$$
   where $H_{\text{KD}}$ is the amino acid's hydrophobicity index.
   - **Timbre (Instrument):** Determined by Zimmerman polarity. A binary mapping is used: non-polar residues are mapped to a Piano sound, while polar residues are mapped to a Cello sound.
   - **Duration:** Proportional to the amino acid's molecular weight, calculated as $MW/100.0$.

2. **Audio Synthesis:** The resulting sequence of musical notes is first encoded into a standard MIDI file using the `midiutil` library. This MIDI file is then rendered into a WAV audio file using the **FluidSynth** software synthesizer with the `GeneralUser GS v1.471` SoundFont.

3. **Harmonic Feature Extraction:** The synthesized WAV file is analyzed using the `librosa` library. The core of the harmonic score is derived from the **Tonal Centroid Features**, also known as **Tonnetz**. The Tonnetz is a representation that projects the audio onto a six-dimensional space derived from music theory, capturing harmonic relationships. The final **Harmonic Score** for a sequence is formally defined as the maximum value within the entire computed Tonnetz matrix:

$$\text{Harmonic Score} = \max(\texttt{librosa.feature.tonnetz}(y, sr)) \tag{6}$$

where $y$ is the audio time series and $sr$ is the sample rate. A higher score indicates a stronger presence of clear tonal centers, which we interpret as a measure of musical ”harmony” or ”consonance”.

#### E.3 Directed Evolution Loop and Multi-Objective Selection

New GFP variants were generated and selected using a multi-step directed evolution algorithm.

1. **Candidate Pool Generation:** A large pool of **1,500** candidate variants was generated. This was done by first randomly selecting parent sequences from the initial training dataset. Each parent sequence was then subjected to point mutations using the `point_mutation_generator` function. The number of mutations introduced into each sequence was not fixed but was sampled from an empirical probability distribution. This distribution was calculated by analyzing the frequency of mutation counts in the original Sarkisyan et al. dataset, aiming to mimic the natural diversity of the experimental variants (Table S5).

Table S5: Empirical probability distribution for the number of point mutations applied per sequence during candidate generation. This distribution was derived from the observed mutation counts in the Sarkisyan et al. (2016) dataset.

| Mutations | Probability | Mutations | Probability | Mutations | Probability |
|---|---|---|---|---|---|
| 2 | 0.000455 | 8 | 0.092924 | 13 | 0.002204 |
| 3 | 0.010684 | 9 | 0.048294 | 14 | 0.000838 |
| 4 | 0.078790 | 10 | 0.026207 | 15 | 0.000168 |
| 5 | 0.307086 | 11 | 0.012241 | 16 | 0.000048 |
| 6 | 0.253809 | 12 | 0.005462 | 18 | 0.000024 |
| 7 | 0.160766 | | | | |

2. **Scoring:** Each of the 1,500 candidates was evaluated using both the trained fitness predictor (to obtain a `predicted_fitness` score) and the harmonic score calculation method (to obtain a `harmonic_score`). This process was parallelized using a `ProcessPoolExecutor` to improve efficiency.

3. **Multi-Objective Selection Strategy:** To select a final set of **300** variants, we employed a two-stage ”supplement” strategy that explicitly balances fitness and harmonic score:

   (a) **Core Fitness Selection:** First, the top **250** candidates with the highest `predicted_fitness` were selected. This forms a ”core elite” set optimized for the primary biological objective.

   (b) **Harmonic Supplementation:** Next, from the remaining 1,250 candidates, the top **50** candidates with the highest `harmonic_score` were selected.

   (c) **Final Combination:** The final set of 300 variants is the union of the 250 fitness-selected variants and the 50 harmonically-selected variants. This approach ensures a baseline of high fitness while using the harmonic score to introduce diverse and potentially interesting candidates that might have been overlooked by a purely fitness-driven selection.

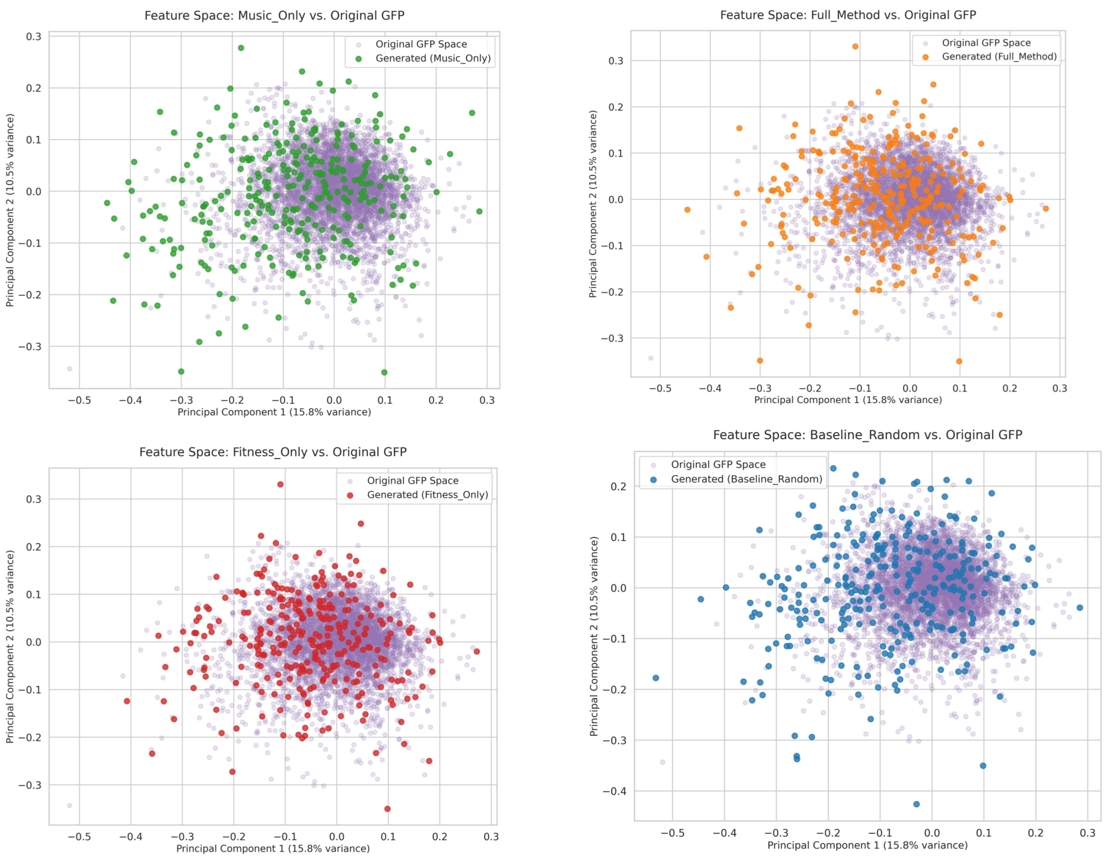

Figure S1: PCA projection of sequences from all four selection strategies. The populations show distinct diversity patterns: `Music_Only` (diversity = **8.1233%**) and `Baseline_Random` (diversity = **7.7387%**) are the most diverse, while `Fitness_Only` (diversity = **7.0261%**) is the most constrained. Our `Full_Method` (diversity = **7.1159%**) achieves a balance, showing greater diversity than the `Fitness_Only` baseline.

### F Case Study: Spectral Feature Analysis of Protein Structures

#### Objective

The purpose of frequency-domain analysis is to characterize the dynamic behavior and structural complexity of protein sequences by analyzing the spectral properties of audio-represented signals. Proteins exhibit unique amino acid compositions, secondary structures, and tertiary structures, which manifest in distinct frequency distributions and spectral features when translated into audio signals. These spectral characteristics facilitate the identification of stable and dynamic regions within protein sequences, highlighting functional variations across different structural domains.

#### Principle

Protein amino acid sequences and structural data can be encoded as audio signals, enabling the use of frequency-domain analysis as a robust tool to extract meaningful patterns from these signals. By calculating features such as Mel-Frequency Cepstral Coefficients (MFCC), spectral centroid, and spectral flatness, we capture essential information about the frequency distribution and dynamic variations within protein sequences. These spectral features not only enhance the understanding of protein structural complexity from an audio perspective but also serve as valuable inputs for classification, prediction, and other computational tasks. This multidimensional analysis of protein structure and function supports further machine learning applications by providing enriched data representations.

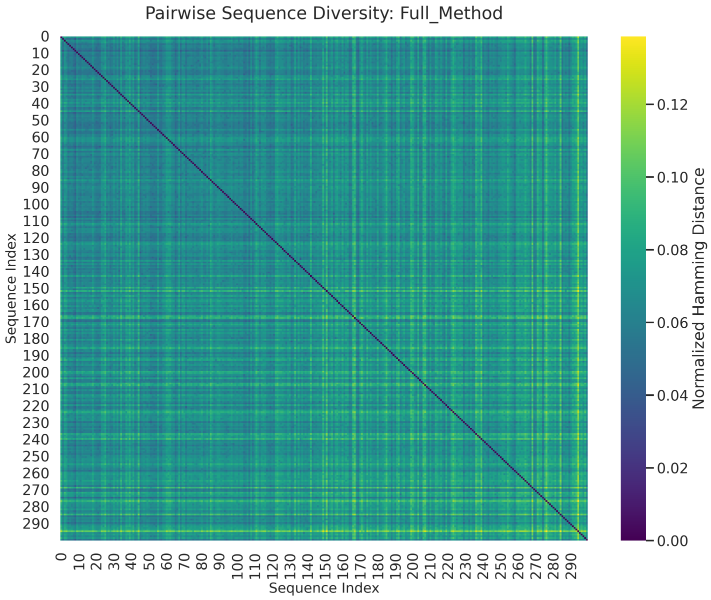

Figure S2: Heatmap of pairwise normalized Hamming distances for a subsample of sequences selected by the `Full_Method`. The color distribution indicates a heterogeneous population with varying degrees of sequence similarity, quantitatively supporting the high diversity score (**7.1159%**).

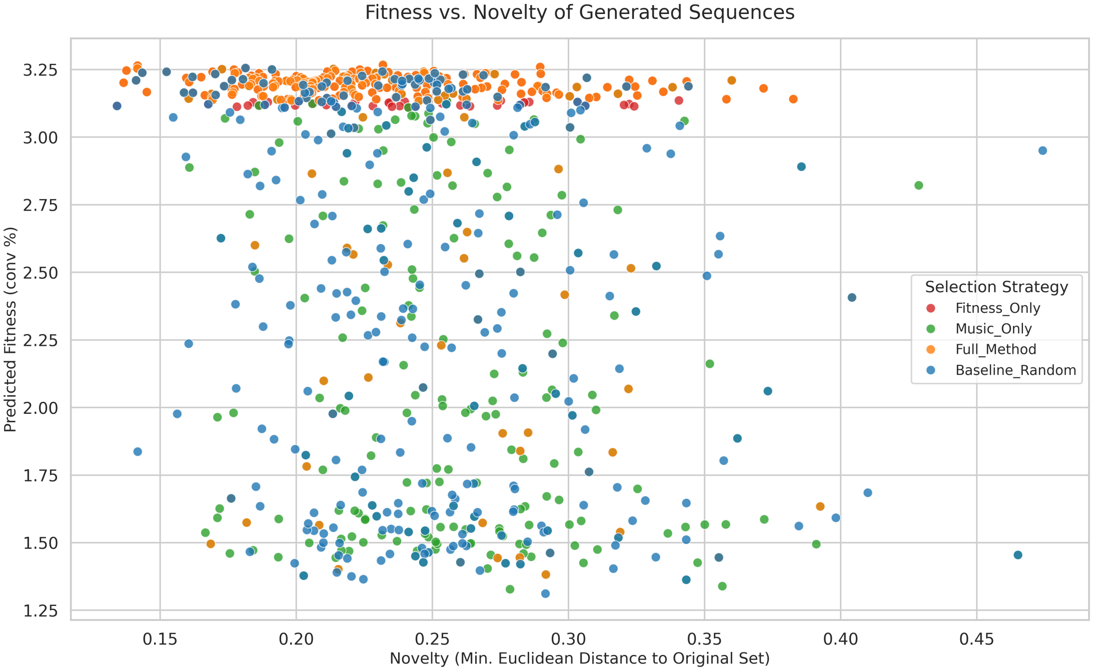

Figure S3: Relationship between predicted fitness and novelty for all generated sequences. Novelty is defined as the minimum Euclidean distance in the ESM-2 embedding space to any sequence in the original dataset. The plot shows that `Full_Method` identifies sequences with both high fitness and a broad range of novelty values.

### Methods

#### F.1 Data Preprocessing and Spectrogram Generation

**Data Loading and Processing.** Audio signals are loaded using the `librosa` library, with the original sampling rate preserved (`sr=None`) to maintain signal integrity.

**Spectrogram Generation.** The Short-Time Fourier Transform (STFT) is performed to generate spectrograms, followed by conversion to a logarithmic amplitude scale (in dB). This transformation highlights the time-frequency variations of audio signals, enabling visualization of structural dynamics and static regions in protein sequences. (See Figure S4)

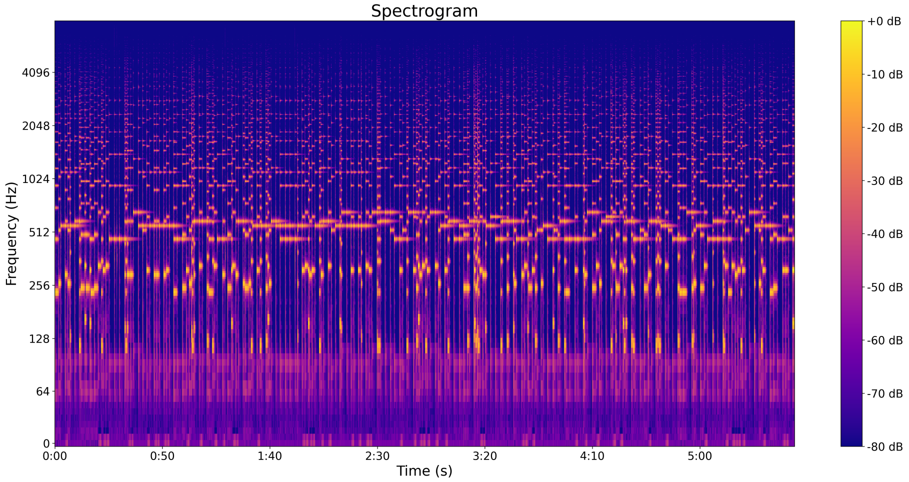

Figure S4: Spectrogram.

**Generation of Log-Mel Spectrogram.** The Mel frequency energy spectrum of the audio signal is computed and converted into a logarithmic scale (dB), revealing patterns in the frequency distribution of the protein audio signal. This approach is particularly effective for identifying functional regions embedded within the signal. (See Figure S5)

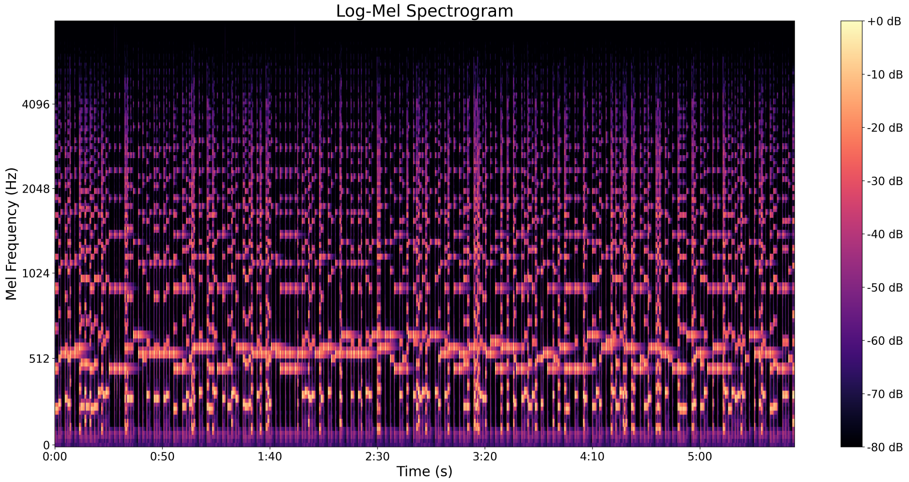

Figure S5: Log-Mel Spectrogram.

**Generation of Power Spectrum.** The power spectrum of the audio signal is computed using STFT to illustrate the temporal changes in frequency. This method highlights the primary frequency components of protein sequences, aiding in the identification of functional regions within the sequence. (See Figure S6)

**Generation of MFCC Image.** Sixty-dimensional Mel-Frequency Cepstral Coefficients (MFCC) are extracted from the audio signal and visualized as a time-varying image. The MFCC image captures the spectral envelope of the audio

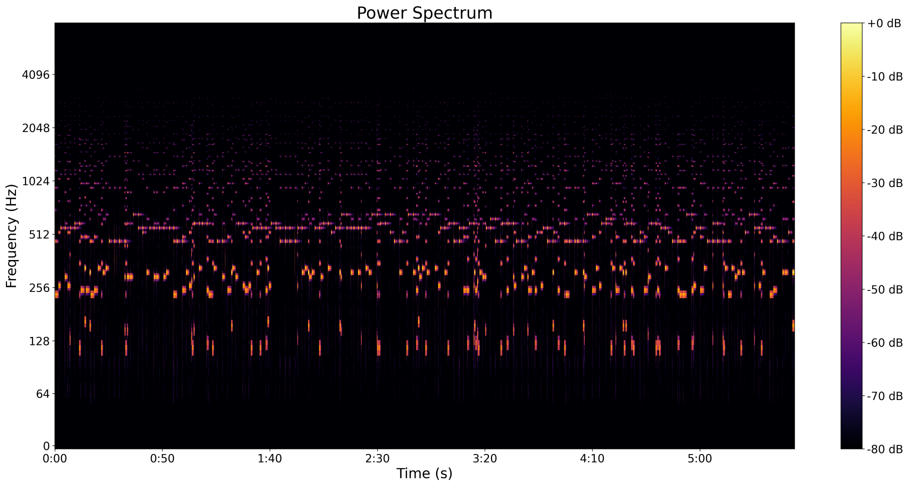

Figure S6: Power Spectrum.

signal by representing its cepstral coefficients, offering insights into the tonal variation of the signal. This approach facilitates the analysis of the overall morphology and complexity of protein sequences. (See Figure S7)

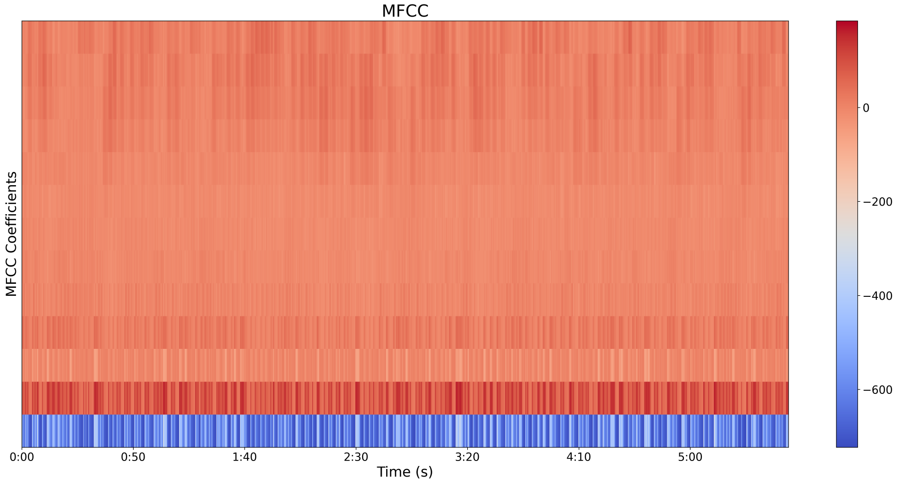

Figure S7: MFCC Image.

#### F.2 Frequency Domain Feature Extraction

To analyze the frequency-domain characteristics of audio signals derived from protein sequences, several key spectral features were extracted, including: Mean MFCC, Spectral Centroid, Spectral Flatness, Spectral Entropy, Zero-Crossing Rate, and Mean Delta MFCC.

#### Results

Four protein sequences from different biological species and functional categories were analyzed: glucose-6-phosphate dehydrogenase (1DPG), $\alpha$-lactalbumin (1A4V), outer membrane lipoprotein (1IWM), and mannose-binding protein A (1AFA). The detailed comparison and analysis of frequency-domain features, considering the amino acid composition and functionality of each protein sequence, are summarized in Table S6.

Table S6: Comparative Analysis of Frequency-Domain Features. Abbreviations: MFCC (Mel-Frequency Cepstral Coefficients), ZCR (Zero-Crossing Rate).

| **Protein** | **Mean MFCC** | **Spectral Centroid (Hz)** | **Spectral Flatness** | **Spectral Entropy** | **ZCR** | **Mean Delta MFCC** |
|---|---|---|---|---|---|---|
| Glucose-6-phosphate Dehydrogenase | -34.28810 | 861.10952 | 0.00007 | 5.60367 | 0.06885 | -0.00087 |
| $\alpha$-Lactalbumin | -33.95147 | 852.43250 | 0.00006 | 1.30511 | 0.06629 | -0.00639 |
| Outer Membrane Lipoprotein | -34.13306 | 872.84715 | 0.00006 | 2.14594 | 0.07173 | -0.00451 |
| Mannose-Binding Lectin-A | -38.04973 | 813.48966 | 0.00005 | 2.53378 | 0.06588 | 0.00126 |

**Mean MFCC.** The mean MFCC of 1AFA is relatively low (-38.05), indicating that its audio signal's spectral envelope is more concentrated in the low-frequency range with a simpler spectral shape, dominated by low-frequency components. In contrast, 1DPG (-34.29) and 1A4V (-33.95) show higher mean MFCC values, suggesting that their spectral envelopes contain more mid-to-high frequency components, indicating greater complexity with higher frequency variability. The multidimensional features of MFCC capture the overall shape of the spectrum, reflecting dynamic changes in the amino acid sequence, especially the distribution of charged or polar amino acids (e.g., lysine K and glutamic acid E), whose diversity leads to a more complex spectral envelope beyond simple frequency centroid shifts.

**Spectral Centroid.** 1IWM has the highest spectral centroid (872.85 Hz), indicating a dominance of high-frequency components in its spectrum, possibly due to its complex structure involved in membrane interactions with the external environment. In contrast, 1AFA has the lowest spectral centroid (813.49 Hz), indicating a predominance of low-frequency components, which may be related to its relatively stable structure.

**Spectral Flatness.** Spectral flatness reflects the smoothness or noise-like quality of the spectrum. 1DPG has the highest spectral flatness (0.00007), indicating a more uniform frequency distribution, potentially suggesting multiple functional regions and dynamic variations. Conversely, 1AFA has the lowest spectral flatness (0.00005), indicating a concentrated frequency component, implying a more stable sequence structure with fewer variations.

**Spectral Entropy.** Spectral entropy measures the complexity of the spectrum, with higher values indicating more complex frequency components. 1DPG has the highest spectral entropy (5.6037), suggesting multiple functional regions in the sequence, leading to higher spectral complexity. In contrast, 1A4V has the lowest spectral entropy (1.3051), indicating a relatively simple spectral structure, consistent with its physiological role as a stable and specialized whey protein.

**Zero-Crossing Rate.** The zero-crossing rate reflects the rate of frequency changes in the signal, with higher rates indicating more high-frequency components and faster frequency variation. 1IWM has the highest zero-crossing rate (0.0717), indicating a rich presence of high-frequency components and rapid frequency variation. Conversely, 1A4V has the lowest zero-crossing rate (0.0663), indicating relatively stable frequency variation, in line with its stable sequence structure.

**Mean Delta MFCC.** The mean Delta MFCC reflects temporal changes in the spectral envelope. 1AFA has the highest mean Delta MFCC (0.00126), suggesting more pronounced spectral envelope fluctuations, which may correspond to dynamic sequence changes when binding carbohydrates. In contrast, 1IWM has a lower mean Delta MFCC (-0.00451), indicating a more stable spectral envelope, consistent with the stable structure of a membrane protein.

This study highlights structural and functional differences among proteins as reflected in their spectral features. Complex proteins like 1DPG exhibit significant complexity in features such as spectral entropy and spectral flatness, while more stable proteins like 1A4V and 1AFA show relatively lower values in these features. The membrane-associated protein 1IWM displays more high-frequency components and rapid frequency changes, reflecting its complex structural domains and dynamic functionality. This detailed analysis of frequency-domain features provides foundational data and new perspectives for further research on protein structure and function.
